# Mechanistic Analysis of Riboswitch Ligand Interactions Provides Insights into Pharmacological Control over Gene Expression

**DOI:** 10.1101/2024.02.23.581746

**Authors:** Shaifaly Parmar, Desta Doro Bume, Colleen Conelly, Robert Boer, Peri R. Prestwood, Zhen Wang, Henning Labuhn, Krishshanthi Sinnadurai, Adeline Feri, Jimmy Ouellet, Philip Homan, Tomoyuki Numata, John S. Schneekloth

**Affiliations:** Chemical Biology Laboratory, Center for Cancer Research, National Cancer Institute, Frederick, MD 21702-1201, USA; Depixus SAS, 3-5 Impasse Reille, 75014 Paris, France; Center for Cancer Research Collaborative Bioinformatics Resource, National Cancer Institute, National Institutes of Health, Bethesda, MD, 20892, USA; Advanced Biomedical Computational Science, Frederick National Laboratory for Cancer Research, Frederick, MD, 21702, USA; Department of Bioscience and Biotechnology, Graduate School of Bioresource and Bioenvironmental Sciences, Kyushu University, Fukuoka 819-0395, Japan

## Abstract

Riboswitches are structured RNA elements that regulate gene expression upon binding to small molecule ligands. Understanding the mechanisms by which small molecules impact riboswitch activity is key to developing potent, selective ligands for these and other RNA targets. We report the structure-informed design of chemically diverse synthetic ligands for PreQ_1_ riboswitches. Multiple X-ray co-crystal structures of synthetic ligands with the *Thermoanaerobacter tengcongensis* (*Tte*)-PreQ_1_ riboswitch confirm a common binding site with the cognate ligand, despite considerable chemical differences among the ligands. Structure probing assays demonstrate that one ligand causes conformational changes similar to PreQ_1_ in six structurally and mechanistically diverse PreQ_1_ riboswitch aptamers. Single-molecule force spectroscopy is used to demonstrate differential modes of riboswitch stabilization by the ligands. Binding of the natural ligand brings about the formation of a persistent, folded pseudoknot structure, whereas a synthetic ligand decreases the rate of unfolding through a kinetic mechanism. Single round transcription termination assays show the biochemical activity of the ligands, while a GFP reporter system reveals compound activity in regulating gene expression in live cells without toxicity. Taken together, this study reveals that diverse small molecules can impact gene expression in live cells by altering conformational changes in RNA structures through distinct mechanisms.

## INTRODUCTION

RNA molecules exert function through specific sequences capable of adopting diverse secondary and tertiary structures.^1, 2^ The formation of dynamic structural ensembles by RNA is governed by a range of factors, including inherent thermodynamic properties as well as contexts and cues within the cellular environment such as protein or small molecule binding.^3^ Within biological contexts, it is frequently the case that an RNA will adopt an ensemble of different but energetically similar three dimensional structures that coexist simultaneously.^4, 5^ Despite this conformational heterogeneity, many RNAs are still capable of folding into structures with hydrophobic pockets that are likely targetable with small molecules.^6, 7^ Small molecules are capable of recognizing these three dimensional pockets by a mechanism that is distinct from sequence-based recognition.^8^ Interactions between RNAs and protein or small molecule ligands can influence the RNA conformational ensembles by stabilization or alteration of populations of structures, leading to modified function.^9^ Changes in structure-dependent regulatory processes are key to normal function, and can lead to the manifestation of various disease states.^10, 11^ Understanding how RNA structures respond to interacting partners is valuable,^12^ as this can be leveraged for drug development as well as a better understanding of endogenous gene regulation with small ligands.^13, 14^

Riboswitches are intriguing systems for investigating mechanistic aspects of RNA-ligand interactions, where well-defined and complex three-dimensional folds have evolved to enable small molecule recognition and altered gene expression.^15, 16^ Riboswitches regulate gene expression specifically through ligand binding to three dimensional folded structures, primarily in bacteria.^17^ One class of commonly identified riboswitches are responsible for sensing the PreQ_1_ metabolite (7-aminomethyl-7-deazaguanine). These riboswitches are associated with regulation of levels of hypermodified guanine, prequeuosine and queuosine itself, a metabolite used in the posttranslational modification of tRNAs.^18^ Evolutionarily diverse PreQ_1_ riboswitches encompass three distinct classes—Class I, II, and III— and differ fundamentally in structure, ligand binding, and gene regulatory mechanisms.^19^ Class I riboswitches feature a singular aptamer domain that directly interacts with PreQ_1_, initiating transcriptional attenuation or termination and in some cases inhibition of translational initiation. ^20, 21^ Class II PreQ_1_ riboswitches possess a distinct architecture, with the aptamer and expression platforms segregated in sequence. This class modulates gene expression through conformational changes upon PreQ_1_ binding.^22, 23^ Notably, Class III riboswitches are characterized by a pseudoknot structure formed by the aptamer domain, enabling ligand recognition and gene regulation.^24^ These variations highlight the multifaceted mechanisms employed by PreQ_1_ riboswitches in bacteria to dynamically regulate gene expression in response to ligand binding. ^25^ Riboswitches have long been considered compelling targets for small molecules. Early studies focused on medicinal chemistry efforts to develop synthetic compounds derived from riboswitches including examples such as FMN^26, 27^, TPP^28, 29^, glmS^30, 31^ PreQ_1_^32, 33^, and lysine^34^. In addition, riboswitches represent rare cases where atomic resolution structures of small molecules in complex with RNA have been solved, enabling detailed biophysical analysis. Previously, our laboratory has studied multiple riboswitches as targets, including the ZMP riboswitch^35^ and multiple PreQ_1_ riboswitches.^36, 37^

Here, we present a structure-informed approach to develop novel, biologically active ligands for PreQ_1_ riboswitches that involves modifying a chemical scaffold to enable biological activity. We report multiple novel synthetic ligands, including **4**, that can directly bind to the PreQ_1_ riboswitch despite no obvious chemical similarity to PreQ_1_ itself. Along with other derivatives, **4** displayed tight binding affinity to the *Bacillus subtilis* (*Bsu*)-PreQ_1_ riboswitch. Investigation of five other PreQ_1_ riboswitches diverse in sequence, structure, function, and evolutionary origin revealed that this synthetic ligand exhibited similar conformational effects to PreQ_1_ in most cases. An X-ray co-crystal structure of the ligand in complex with a PreQ_1_ riboswitch revealed an identical binding site, but distinct binding mode relative to PreQ_1_. A second new scaffold, based on the harmol heterocycle (**8**), was also co-crystallized with the aptamer, and evaluated in functional assays. This compound displayed *in vitro* activity but was inactive in cell-based assays, potentially due to the more promiscuous nature of the scaffold interacting with other RNAs. Single molecule assays revealed that PreQ_1_ induces stable pseudoknot formation. However, the binding of **4** impacts riboswitch function by a distinct kinetic mechanism, altering the rate of folding and most likely stabilizing the PreQ_1_ RNA in a partially folded “pre-pseudoknot” state, despite having the same conformational consequence in bulk measurements. Both *in vitro* transcription termination and *in vivo* expression assays in bacterial cells validate the ability of **4** to impact gene expression by binding directly to RNA. This work demonstrates that diverse chemical scaffolds can bind to and influence riboswitch aptamers to accomplish similar functional outcomes through distinct mechanisms.

## RESULTS AND DISCUSSION

### Structure-informed alterations in chemical structure impact binding to aptamer

Previous work demonstrated that a synthetic dibenzofuran ligand has high affinity and selectivity to both *Bsu*-PreQ_1_ and *Tte*-PreQ_1_ riboswitches.^36^ We used ICM MolSoft software^38^ to dock various chemical scaffolds related to the initial dibenzofuran hit compound that had been reported previously. We conducted structural modifications and synthesized various xanthone derivatives to assess their biologically relevant interactions with the PreQ_1_ riboswitch. By altering the side chains and incorporating other modifications, as detailed in Table 1, we synthesized several analogs and examined their recognition ability and activity. A goal of this exercise was to identify new, synthetically accessible chemical scaffolds potentially capable of improved affinity or activity by making more contacts to the RNA. Using this approach, we designed and synthesized nine small molecule ligands representing new heterocyclic cores or sidechains that could plausibly bind to the PreQ_1_ aptamer (Figure S6).

To evaluate the binding of each compound to the RNA, we employed fluorescence titrations or microscale thermophoresis (MST).^39^ Compounds **1, 2, 4, 5**, and **8** showed changes in ligand fluorescence with increasing concentrations of RNA. For these compounds, ligand fluorescence was plotted as a function of RNA concentration. Data were fitted using a one-site total binding model to measure an equilibrium dissociation constant (K_D_) for each of these compounds. Next, compounds that did not show any fluctuation of ligand fluorescence, (**3, 6, 7**, and **9**) were evaluated using MST using a Cy5-labeled *Bsu* PreQ_1_ aptamer. By fitting the curves as a function of ligand concentration and using a one-site total binding model, an equilibrium dissociation constant (K_D_) was measured (Figure 1A, Figure S1). In general, most of the ligands bound to the RNA with low micromolar affinity. However, compound **9** showed no binding up to a concentration of 500 μM. Of the remaining compounds, **4** showed micromolar binding (K_D_= 16 ± 21 μM), and therefore binding was also evaluated with another dye, AlexaFluor 647, to rule out potential effects due to the fluorophore used in binding analysis. Using labeled aptamers from *Staphylococcus saprophyticus* (*Ssa*)*-*PreQ_1_ and *Tte-*PreQ_1_ that have a conserved binding domain, compound **4** demonstrated apparent K_D_ values of 21.9□±□2.25□μM and 29.0□±□2.4□μM, respectively, in MST (Figure S2), confirming direct binding to the RNA. Next, compounds were used in *in vitro* assays for functional evaluation.

**Figure 1:**
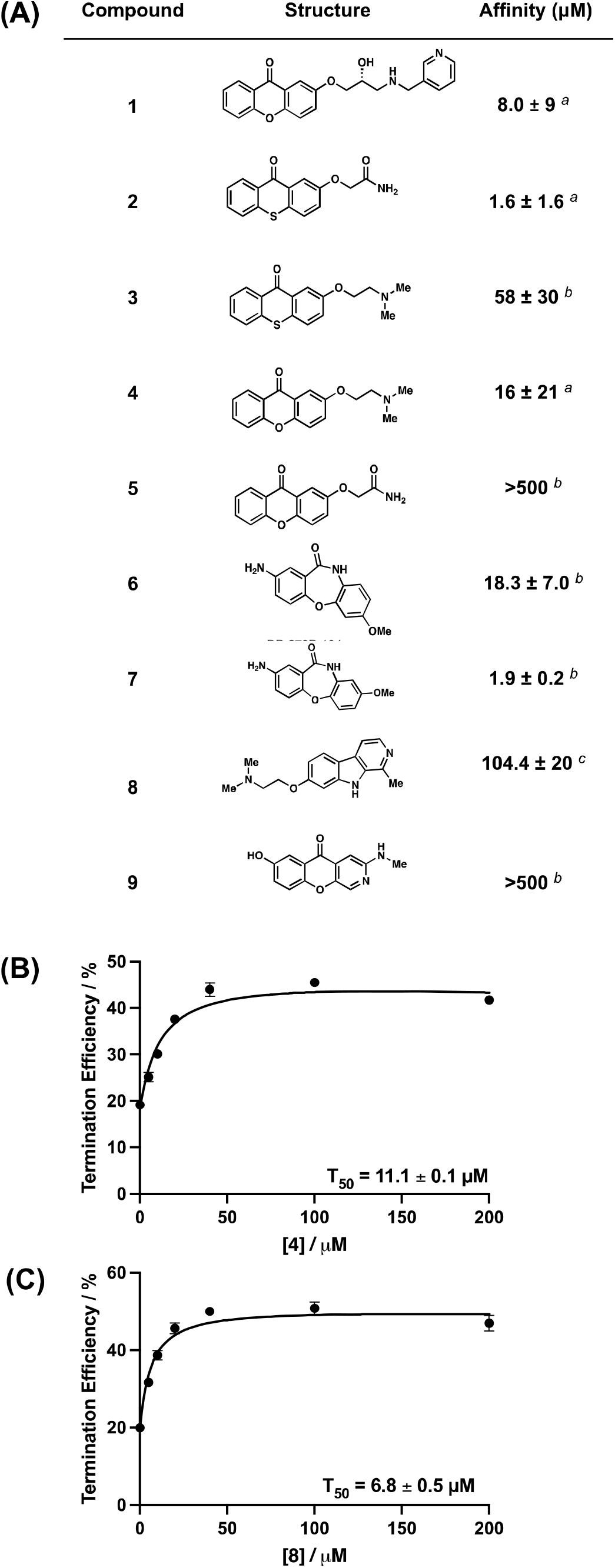
(A) Chemical structures of PreQ_1_ riboswitch aptamer ligands and their binding affinities to a *Bsu*-PreQ_1_ aptamer. ^*a*^ is K_D_ measured using intrinsic ligand fluorescence, ^*b*^ is K_D_ measured by MST measurements, ^*c*^ is K_D_ measured by FIA. (B, C) Quantification of transcription termination efficiencies (T_50_ values) of **4** and **8**, respectively, as a function of concentration of each of the ligand in single round transcription termination assays. Error values represent standard deviation from three independent replicates.

### Synthetic ligands are active in single round transcriptional termination assays

*In vitro* single round transcription termination assays were performed to biochemically analyze the activity of analogs in functional assays. The *Ssa*-PreQ_1_ riboswitch was subjected to transcription in the presence of increasing concentrations of ligands, which could either result into a full transcript read through (RT) or a transcription termination due to the formation of terminator hairpin (T). T and RT products were visualized on a denaturing PAGE and termination efficiency (T_50_) value was calculated by dividing the terminated band intensity by the total RNA intensity. These data show enhanced biochemical activity of **4** (T_50_= 11.1 ± 0.10 μM) and **8** (T_50_= 6.8 ± 0.45 μM) *in-vitro* relative to the initial dibenzofuran. (Figure 1B, C, Figure S3). For the remaining compounds, saturation was not observed at the limit of solubility, and therefore accurate T_50_ values could not be measured. Since compounds **4** and **8** showed activity in functional assays, they were studied further.

### X-ray co-crystal structure establishes ligand binding mode

To further understand the binding mode of **4** and **8**, we performed X-ray crystallography on the ligand aptamer complex. The co-crystal structures of the abasic mutant at positions 13, 14, and 15 in *Tte-*PreQ_1_ riboswitch aptamer (ab13_14_15) with **4** and **8** were determined at 2.15 A□ and 2.25 A□ resolution, respectively, by molecular replacement method (Figure 2A, 2B and Table S4). Compound **4** binds at the PreQ_1_ binding site, where the xanthone core is sandwiched by one face with G11 and the other with G5 and C16, residues that are strictly conserved in the class I PreQ_1_ riboswitches. When the current co-crystal structure is superimposed onto the PreQ_1_-bound form, the planar rings of their ligands are well overlapped (Figure 2C). However, because **4** is bulkier than PreQ_1_ and its heteroatom content is less than that of PreQ_1_, the binding pose of **4** slightly diverges from that of PreQ_1_. In the co-crystal structure with PreQ_1_, one side of the base containing the N2, N3, and N9 atoms of PreQ_1_ is recognized by strict hydrogen bonds with the N1 and N6 atoms of A29 and the O4 atom of U6 of the *Tte*-PreQ_1_ riboswitch, respectively (Figure 2D). In contrast, the corresponding side of the xanthone moiety of **4** is further from these crucial atoms, resulting in a tilted binding axis of the heterocyclic core of **4** compared to PreQ_1_ of approximately 15 degrees (Figure 2C). Consequently, the oxygen atom of the central ring of **4** is situated at 3.6 A□ away from the N6 atom of the phylogenetically conserved A29. This finding suggests a weak hydrogen bonding interaction between the riboswitch and compound **4**, unlike the strong interaction observed in the PreQ_1_-bound form where the distance between the N6 atom of A29 and the N3 atom of PreQ_1_, a counterpart of the oxygen atom of the central pyranoid ring of **4**, is 3.1 A□. Superimposition of these two structures indicates that **4** collides with the base of C15 of the PreQ_1_-bound structure, due to the size of **4** being larger than that of PreQ_1_. Consequently, the conformation of the sugar and phosphate backbone at position 15 of the current structure is relocated to fit **4** into the ligand binding site, when compared to the PreQ_1_-bound form. It is important to note that C15 of the riboswitch is critical for recognizing PreQ_1_ via the canonical Watson-Crick base pairing. Therefore, the binding of **4** to the PreQ_1_ binding site would affect the conformations of L2 and S3 that are important for regulating the riboswitch function. Consistent with this, the co-crystal structure with **4** exhibits conformational differences of L2 and S3 when compared to those in the PreQ_1_-bound form. Since the abasic mutant at positions 13, 14, and 15 was used in this study, we cannot rule out the possibility that the conformational differences are due to the introduction of the abasic sites in the current construct. However, given the steric hindrance between **4** and C15, **4** probably has a major effect on the structure of these regions, which could be related to the differences in the results of biochemical analyses described below.

**Figure 2:**
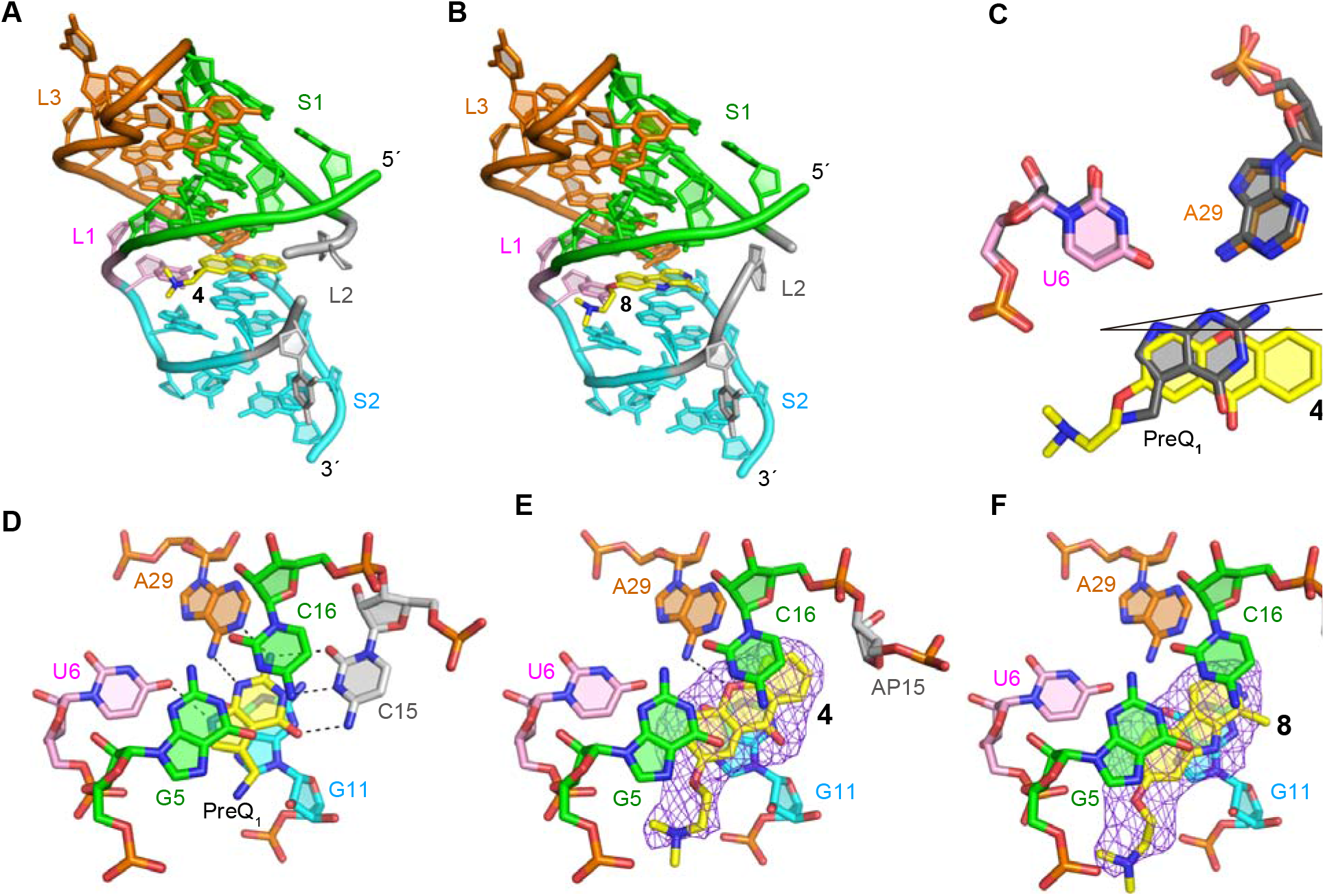
X-ray crystal structures of ab13_14_15 in complex with synthetic ligands. (A,B) Overall structure of the complexes with ligands **4** (A) and **8** (middle). (C) Comparison of binding poses between **4** and PreQ_1_. (D) Structural comparison between the wild-type *Tte*-PreQ_1_ riboswitch aptamer complexed with PreQ_1_ (PDB ID: 3Q50), (E) ab13_14_15 complexed with **4** and (F) ab13_14_15 in complex with **8**. Hydrogen bonds are shown in dashed lines. Purple mesh represents the m*F*_o_-D*F*_c_ electron density maps observed for each ligand, which are contoured at 2.5 σ. The distance between the oxygen atom of the central ring of **4** and N6 atom of A29 is 3.6 A□.

Like **4**, compound **8** is situated in the PreQ_1_ binding site and is surrounded by the phylogenetically conserved nucleotides. While **4** forms a hydrogen bond with the riboswitch, **8** does not hydrogen bond with any nucleotides. Therefore, **8** is stabilized in the ligand binding site of the riboswitch by stacking and hydrophobic interactions. Compared to the binding site of **4**, the binding site of **8** is shifted about 1-2 A□ in the opposite direction of the L2 loop. This is likely because **8** has no hydrogen bond with the riboswitch, which anchors the compound in the ligand binding site (Figure 2F). In our previous report^36^, we analyzed the effects of dibenzofuran and carbazole derivatives on PreQ_1_ riboswitch function and showed that the binding poses of these compounds differ due to changes in the acceptor/donor pair of hydrogen-bond between these compounds and the riboswitch. Ligand **8** is a derivative of harmol and has a nitrogen atom in the central ring like the previous carbazole derivative. However, the binding pose of **8** is quite different when compared other nitrogenous heterocycle ligands (such as PDB ID: 6E1V), and the nitrogen atom of the central ring of **8** faces in the opposite direction. Therefore, the conserved nucleotides, U6 and A29, that are crucial for recognizing PreQ_1_ by hydrogen-bonds only contact with the heterocycle of **8** via van der Waals interactions. Together, these structures provide a rationale for both how diverse ligands recognize the aptamer binding site and why they are active in functional assays.

### Structure probing reveals impacts of ligand binding on aptamer flexibility

PreQ_1_ riboswitches are among the most commonly evolved riboswitches, and as such, have been observed to have considerable diversity in terms of sequence, structure, and mechanisms. Given the diversity of RNA structures that recognize the PreQ_1_ metabolite, we asked whether evolutionarily diverse PreQ_1_ aptamers have differential effects on ligand-mediated recognition and flexibility. We utilized selective 2’-hydroxyl acylation analyzed by primer extension and mutational profiling (SHAPE-MaP) to assess the flexibility of bases at single nucleotide level in the presence and absence of both PreQ_1_ and **4**.^40^ PreQ_1_ RNA aptamers from six different species were selected, representing all three classes of PreQ_1_ riboswitch. We studied aptamers from *Tte*^19, 41^, *Bsu*^19, 41^, *Faecalibacterium prausnitzii (Fpr)*^24^, *Ssa*^36^, *Lactobacillus rhamnosus (Lrh)*^41-43^, *Streptococcus pneumoniae (Spn)*.^*44, 45*^

Each *in-vitro* synthesized RNA was folded and incubated with DMSO, PreQ_1_ or **4**, followed by incubation with the SHAPE reagent 2A3.^46^ Modified and unmodified RNAs were reverse transcribed, and mutations were mapped by next generation sequencing. Data analysis using the Shapemapper pipeline^47^ revealed mutation rates and the reactivity profile for each nucleotide. Here, lower SHAPE reactivity depicts decreased flexibility (or stabilization) of each nucleoside in the presence of ligand. Next, SHAPE constraints for each nucleotide were utilized to predict secondary structure with RiboSketch software (Figure 3A-F).^48^ Base-pairing probabilities using the SHAPE derived data are shown using the arc plots using Superfold^49^. Delta SHAPE analysis was then used to identify nucleotides specifically altered in flexibility upon binding to the ligand.^50^ After accessing significant changes in the SHAPE reactivity within different riboswitches, *Bsu, Lrh, Spn, Fpr* showed alteration of structure in presence of both ligands. Specifically, the *Bsu* riboswitch showed stabilization of structure at C20, A21, C22 belonging to aptamer domain, (consistent with crystallographic studies),^18^ as well as at the C53 and U60 bases within the terminator domain. However, A42, C43, G44 and terminator hairpin bases U55, U56, G57 display destabilization in presence of PreQ_1_ ligand (Figure S4A). With PreQ_1_ and the *Spn* aptamer, A52, G53, G54, A55, G56 (belonging to the loop J2-4) were stabilized and A41, U42, A43, A44, C45 (that makeup the P4 stem)^44^ are destabilized. This effect was strikingly similar in the presence of **4**, as A41, U42, A43 were stabilized along with the destabilization of A41, U42, A43, A44 bases (Figure S4B,C). The *Lrh* aptamer in presence of PreQ_1_, A31, U32, U33, C36, U37, U38 (J2-3 loop), G57 (P4 region) were observed to have positive delta SHAPE inferring stabilization and bases U49, A50, U 51, U52, A53 (J2-4 loop), A59, A60 (P4 region) had negative delta SHAPE (informed from crystal structure).^51^ With **4**, the effect was similar to PreQ_1_, where bases U30, A31, U32, U33, C36, U37 along with U49, A50 displayed stabilization in the aptamer. However, bases G40, A41, U42 (P3), U52, A53 (J2-4), A69, G70, G71, A72, incorporated in the ribosome binding site (RBS) showed significant destabilization (Figure S4D,E). In *Fpr* PreQ_1_-riboswitch, with ΔSHAPE, only destabilizing events were captured with both ligands, as bases G110, G111, A112, G113 (constituting ribosome binding site) had enhanced reactivities. With PreQ_1_, only A115 showed positive ΔSHAPE, meaning stabilization (Figure S4F,G). In contrast, the *Tte* PreQ_1_ riboswitch displayed only destabilization in the presence of PreQ_1_ (at A24, C25, A26, A27, A28, A29, which have no interaction with the PreQ_1_ ligand) and had no significant ΔSHAPE reactivity when **4** was bound (Figure S4H). In general, all structures displayed decreased reactivity in the presence of both ligands in comparison to the DMSO control, reflecting an overall stabilization of the structure. Both natural (PreQ_1_) and synthetic ligand (**4**) had strikingly similar effects on RNA conformation except for a few nucleotides. While these secondary structures are experimentally informed, they do not necessarily reflect three-dimensional aptamer structure with perfect accuracy. Still, this comparative analysis is a powerful demonstration that chemically distinct ligands can have similar effects on structurally diverse RNAs that recognize a common cognate ligand.

**Figure 3:**
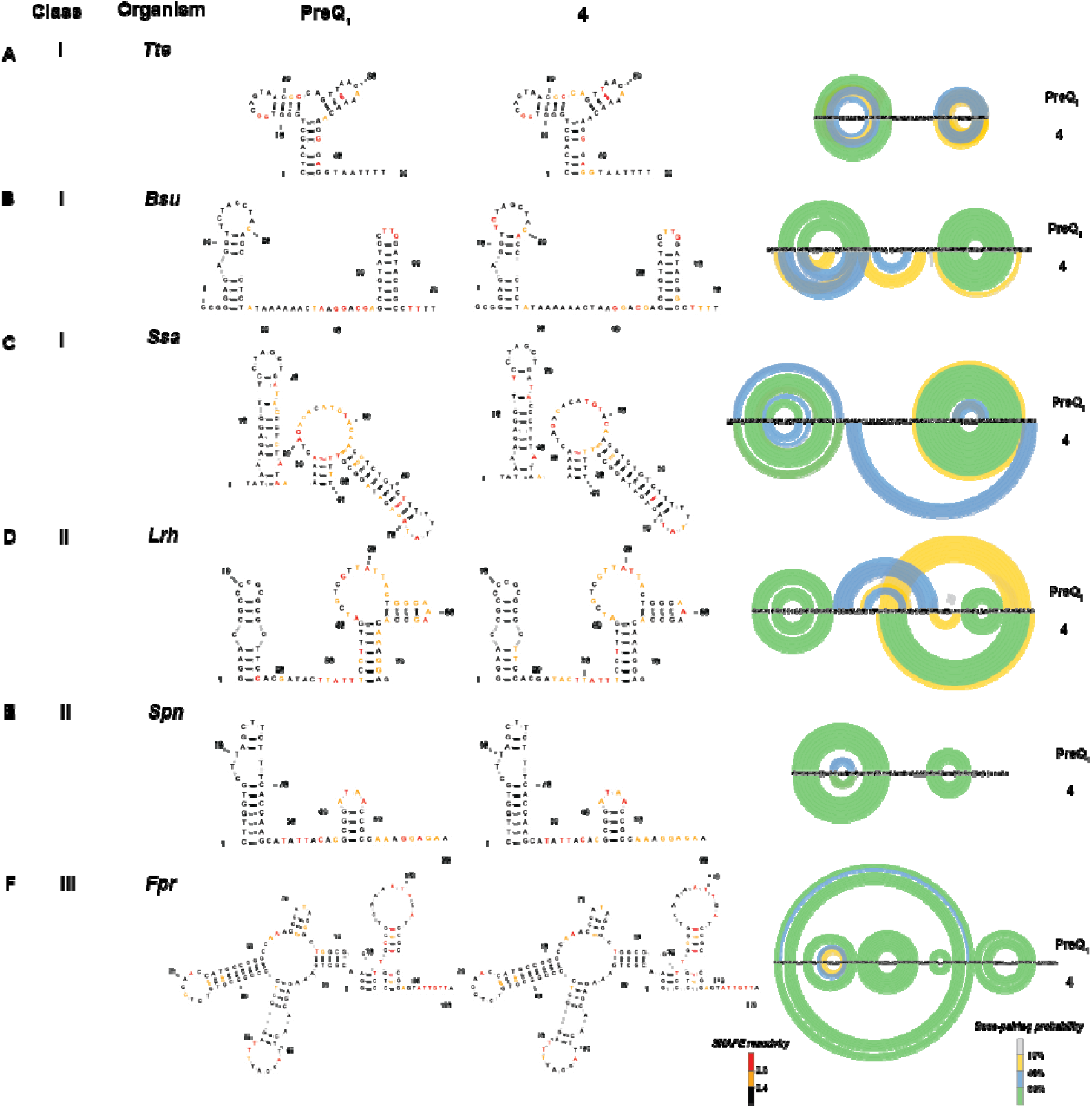
(A-F) SHAPE-MaP informed secondary structure predictions of various riboswitches in the presence of cognate (PreQ_1_) and synthetic ligand (**4**). Arc plots show the SHAPE-MaP profiles of each riboswitch in the presence of PreQ_1_ and **4**. SHAPE reactivity and base pairing probabilities are indicated using the respective color schemes shown at the bottom.

### Ligand binding stabilizes RNA structure

Having confirmed that ligand binding leads to significant changes in the structure of PreQ_1_ RNA using bulk methods in solution, we next used the MAGNA magnetic force spectroscopy platform to evaluate the effects of ligand binding on the stability of the aptamer’s structure at the single molecule level. This platform allows precise tracking of molecular extension in response to an applied force across hundreds of single molecules in parallel to gain insights into molecular dynamics and interactions. To use MAGNA, first, a biotinylated *Bsu*-PreQ_1_ aptamer was bound to a streptavidin paramagnetic bead and tethered to a flow cell floor via hybridization to a surface-bound oligonucleotide. A precisely controllable magnetic force was then applied to the beads whilst their vertical or Z-positions were tracked in real time. When the RNA was subjected to low force, it folded freely. As the force was increased, structural disruption or unfolding occurred, resulting in a sudden change in vertical bead position. The force could then be reduced, allowing the structures to return to a folded conformation (Figure 4A). This non-destructive process was repeated over multiple cycles of slowly increasing, then decreasing forces (referred to as force ramp experiments, Figure S5A), while the forces at which individual structures unfolded and refolded were measured. Addition of ligands to the flow cell allowed tracking of their impact on the stability of the RNA structures, through their effect on these unfolding and folding forces. Separately, stepped constant-force experiments were performed where RNA molecules were subjected to the same force for a fixed amount of time before increasing the force in a stepwise manner (Figure S5A). During each force step, the transition of the RNA between the unfolded and folded states was tracked, and the time spent in the unfolded state was observed to increase with force until the RNA structures remained constantly unfolded. The equilibrium force at which the RNA spent equal time in each state was also determined. Constant force experiments in which the RNA was subjected to the equilibrium force for an extended period could then be performed, to allow the impact of ligand binding on folding and unfolding dynamics to be explored through changes in the equilibrium force and/or the frequency of folding-unfolding events (Figure S5A).

**Figure 4:**
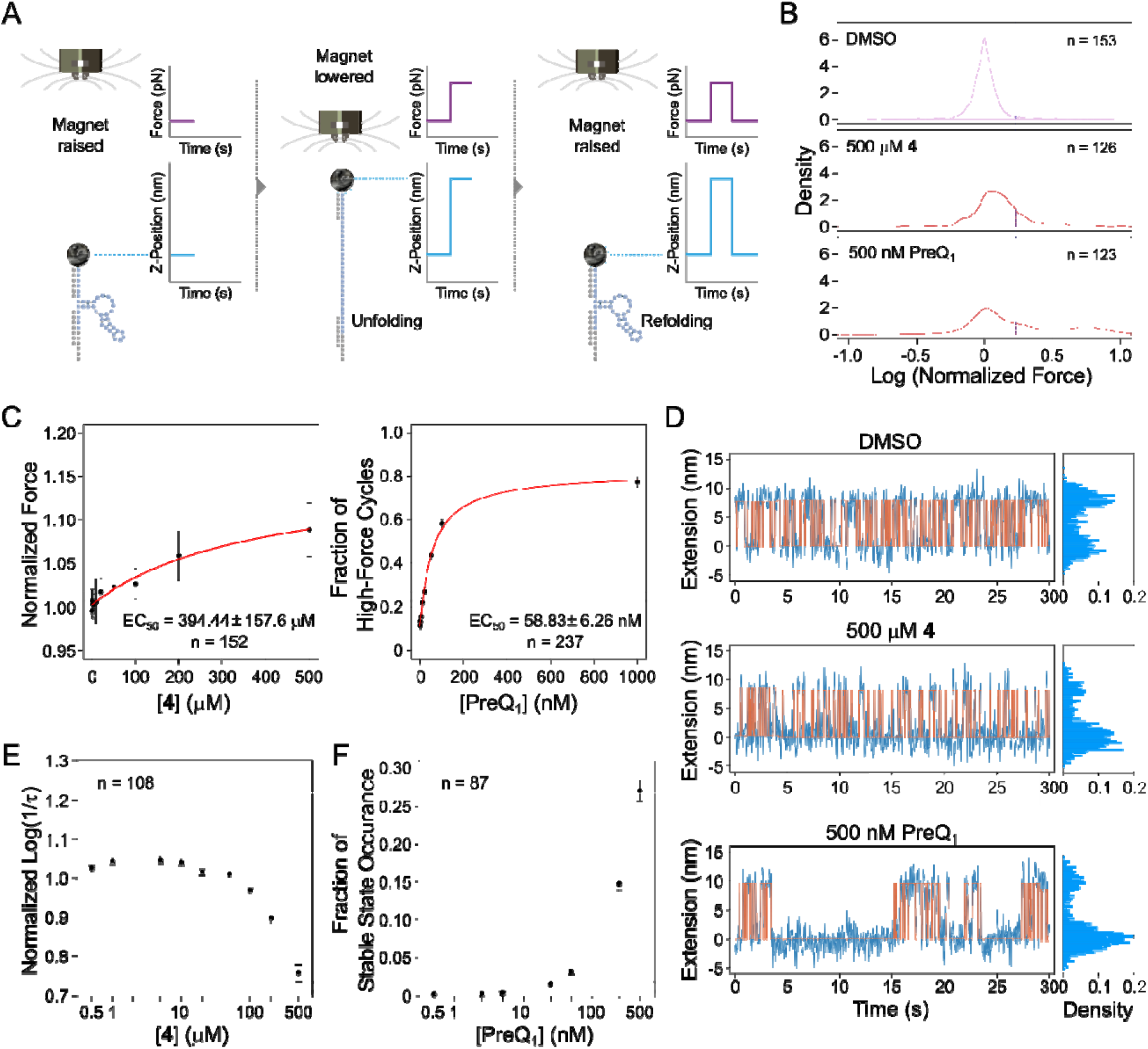
(A) Overview of MAGNA as a single-molecule platform for exploring the interactions of bioactive small molecule ligands with their target RNA structures in real-time (B) Unfolding force distributions of the *Bsu* PreQ_1_ riboswitch aptamer in control, **4** and PreQ_1_ ligand conditions (C) Dose-response curve for the change in unfolding of the aptamer in the presence of **4** and PreQ_1_ (D) Raw traces of the constant-force experiments for control, **4** and PreQ_1_ of a single molecule with cumulative density histograms shown to the right. (E) The aptamer unfolding rate as a function of the concentration of **4** in constant force experiments. (F) The impact of PreQ_1_ ligand concentration on the occurrence probability of the stable folded state. n is the number of molecules analyzed.

We conducted ramp experiments to probe RNA structure unfolding under varied conditions: control (1% DMSO), **4**, and PreQ_1_ and plotted the distribution of the normalized forces required to unfold and refold the RNA structures. In the control condition, the force distribution formed a single peak (Figure 4B) which was attributed to unfolding of a “pre-pseudoknot” structure. Introduction of a saturating concentration (500 μM) of **4** subtly shifted the peak of the force distribution toward higher forces, implying minor structural influence that increases the force needed to unfold and refold the structure (Figure 4B & Figure S5B). In contrast, saturating concentrations of PreQ_1_ (500nM) induced a second peak in higher forces, indicating that the molecules sometimes required a much higher force to unfold, which was attributed to the formation of stable pseudoknot structures. However, the position of the first peak did not shift, demonstrating that PreQ_1_ did not change the stability of the pre-pseudoknot structure (Figure 4B and Figure S5B. The second high force peak was notably absent with **4**, highlighting that the compound did not trigger the formation of persistent/stable pseudoknots like those induced by PreQ_1_ (Figure 4B).

The increase caused by compound **4** on both the median normalized unfolding and refolding forces was shown to be concentration dependent (Figure 4C and Figure S5C respectively) with EC_50_ values of 394±157 μM and 456±258 μM respectively, suggesting that **4** stabilizes the riboswitch pre-pseudoknot structure and that this interaction helps to refold the RNA. For the PreQ_1_ ligand, concentration effect was assessed differently, using the fraction of high force cycles, to account for cycles in which the RNA in pseudoknot conformation did not unfold at the maximal force applied. PreQ_1_ binding increased the fraction of the high-force cycles in a concentration dependent manner indicating an increase in pseudoknot formation whilst RNA refolding was not affected by the cognate ligand (Figure 4C and S5D).

Under constant-force experiments, the bead position tracking of individual molecules showed that the PreQ_1_ ligand prolonged folded state duration compared to the control and **4** (at the same applied forces), revealing ligand-induced pseudoknot formation (Figure 4D, S5E). Analysis of the lifetimes of the folded states under the control, PreQ_1_ (500 nM) and **4** (500 μM) conditions showed an exponential distribution of the observed events with **4** inducing a slightly increased lifetime compared to the control. In the presence of the cognate ligand, the folded states fitted a combination of two distinct lifetime distributions (confirmed using the Bayesian information criterion) (Figure S5F, representing one single molecule). Of these two lifetimes, the shorter of the two showed a lifetime similar to that of the control, most likely corresponding to the pre-pseudoknot state, and the second corresponded to the stable folded state attributed to the pseudoknot.

To evaluate concentration dependency of the effects, lifetime data from multiple molecules were aggregated by evaluating log (1/lifetime) and normalizing each condition to the control for the same molecule before combining data from multiple molecules. Compound **4** was confirmed to decrease the unfolding rate (i.e., to cause the RNA to stay longer in the folded pre-pseudoknot state) in a concentration dependent manner, but only at concentrations above 100 μM (Figure 4E). In contrast, the cognate ligand did not affect the unfolding rate of either the short lifetime form (pre-pseudoknot) or long lifetime form (pseudoknot) (Figure S5G) and no change in the refolding rate (Figure S5H). Instead, the probability of the pseudoknot state occurring increased with PreQ_1_ ligand concentration (Figure 4F), confirming that binding of the PreQ_1_ ligand induces pseudoknot formation in a concentration-dependent manner. Compound **4** was also demonstrated to increase the refolding rate in a concentration dependent manner above 100 μM, suggesting that the molecule alters the rate of refolding of the RNA, perhaps by binding to a less folded form (Figure S5I). However, while the PreQ_1_ ligand affects the rate of pseudoknot formation, it has no effect on the less stable pre-pseudoknot structure’s folding dynamics.

### Ligands affect riboswitch activity in cells

Next, we evaluated the ability of **4** to modulate riboswitch activity *in vivo*. We employed an engineered green fluorescent protein (GFPuv) reporter assay, as has been used previously to demonstrate riboswitch activity.^52,53^ The *Bsu-* PreQ_1_ riboswitch aptamer was cloned into a plasmid bearing GFPuv expression in parallel with a second, empty vector that expresses GFP but lacks any riboswitch. Next, GFPuv positive constructs were transformed into the JW2765 strain of *E*.*coli* bearing a Δ*queF* mutation ^54, 55^ to generate a stable cell line expressing the reporter construct. The Δ*queF* mutation leads to impaired PreQ_1_ biosynthesis and provides an ideal system to study the effects of ligands on the PreQ_1_ riboswitch, as it lacks endogenous PreQ_1_. Next, cells were grown on specialized CSB media (to further hinder any endogenous PreQ_1_ biosynthesis) in the presence of compounds or the DMSO control. When visualized under UV, cells grown in the presence of DMSO exhibited high levels of fluorescence (Figure 5A). In contrast, treatment with PreQ_1_ and **4** led to a complete loss of fluorescence levels (Figure 5B, 5C). The cell line expressing an empty vector was not responsive to ligand (Figure 5D). In addition, a compound structurally similar to **4** that did not bind to the riboswitch (compound **9**) was also inactive. Finally, cells treated with **8** also did not respond, even though **8** both binds to and modulates the function of the riboswitch *in vitro*. Importantly, these results demonstrate that both PreQ_1_ and **4** clearly exhibit gene modulation activity by directly binding to RNA structures in cells, rather than nonspecific or other off-target mechanisms.

**Figure 5:**
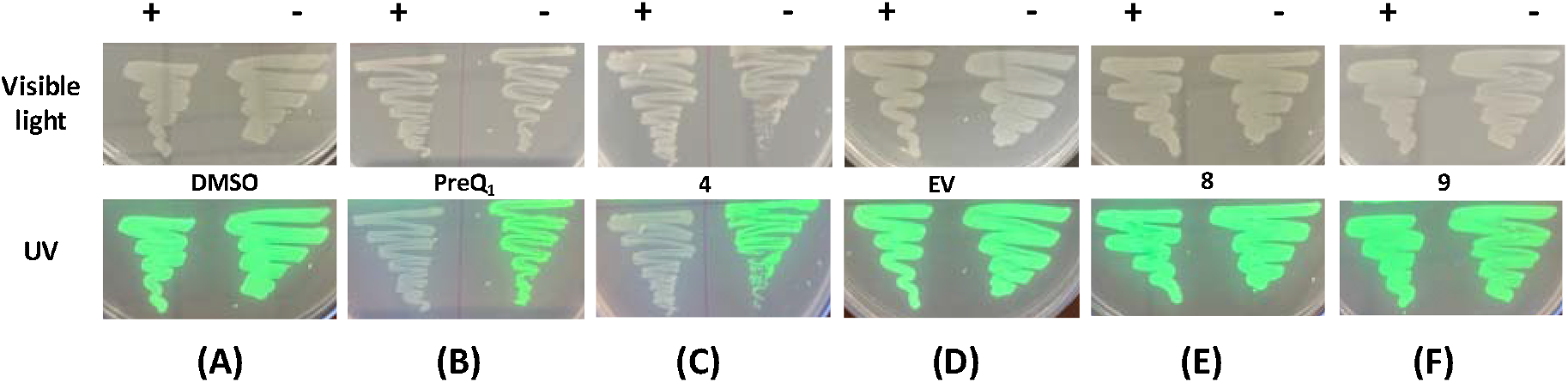
Ligands impact expression of a GFP reporter gene containing a PreQ_1_ aptamer in mutant *E*.*coli* grown on specialized media in the presence and absence of: DMSO (A), PreQ_1_ (B), **4** (C, D), **8** (E) and **9** as negative control (F) visualized under visible light (top) and UV transilluminator (bottom). EV: empty vector.

## CONCLUSIONS

In this work, we demonstrate that structure informed design can be used to identify novel small molecules that bind to RNA structures and impact their function in biophysical, biochemical, and biological assays. By investigating diverse heterocyclic scaffolds, we identified new compounds with considerably improved activity in single round transcription termination assays. Here, subtle changes in compound structure can have dramatic impacts on both binding affinity and activity in functional assays, resulting in a complex and non-obvious structure-activity relationship (SAR). X-Ray crystallographic analyses indicate that multiple new scaffolds can bind to the PreQ_1_ aptamer RNA at the same binding site as the cognate ligand with well-defined binding modes. Further, some ligands exhibited additional specific bonding interactions with the RNA.

In addition to binding assays, SHAPE-MaP studies were used to observe ligand-induced effects on the stability of six evolutionarily and functionally diverse PreQ_1_ riboswitches, all of which recognize the same cognate ligand. In each instance, the synthetic ligand exhibited effects similar to the native ligand. This was demonstrated through similar alterations in base-pairing interactions contributing to the stabilization of the riboswitch structure despite lower binding affinity of the synthetic ligand. Thus, although these RNAs represent distinct sequences, are structurally diverse, and exhibit switching activity through different mechanisms, they recognize small molecules through conserved three-dimensional structures rather than sequence.

Additionally, single molecule magnetic force spectroscopy was used to assess the ability of PreQ_1_ and **4** to impact aptamer folding. Here, the presence of cognate ligand led to the formation of a stable pseudoknot structure. In contrast, the binding of **4** appears to have a distinct mechanism. These experiments indicate that **4** stabilizes the PreQ_1_ RNA, most likely in the pre-folded pseudoknot state. Thus, **4** has a different impact on the folding pathway compared with the cognate PreQ_1_ ligand, impacting the rate of re-folding rather than inducing a stable pseudoknot formation. More specifically, activity observed by the recognition of **4** is possibly due to kinetic alterations rather than thermodynamic factors as with PreQ_1_. Riboswitches are known to exhibit diverse folding pathways and mechanisms of switching, reflecting a complex interplay between ligand binding and structural dynamics that results in altered gene expression. Although in many cases persistent formation of a folded state is thought to be required, it has also been observed that alteration of folding kinetics can drive switching behavior as seen with the ZTP riboswitch.^56,57^

Finally, we used a fluorescent reporter assay to demonstrate that **4** is cell permeable and modulates riboswitch activity in live cells by directly interacting with the RNA. Importantly, both empty vector and a chemically related, non-binding control compound showed no activity. Interestingly, the harmol-based ligand **8** displays weak binding but stronger inhibition in functional biochemical assays and was inactive in the *in vivo* reporter assay. While the reason for the lack of activity is unclear, it may reflect more promiscuous binding of harmol-like molecules to diverse RNA structures, which would presumably require a higher concentration to observe activity given the higher abundance of other RNAs in cells. Alternately, activity in the biochemical assay could be occurring due to nonspecific RNA binding. For example, related β-carboline alkaloids such as harmine and harmaline engage with RNA bases through interactions involving the O^2^ of cytosine and uracil, the N^7^ of guanine and adenine, as well as the phosphate group in the backbone, and can engage in intercalative interactions of diverse RNAs^58^. Thus, not only binding affinity and mechanism of recognition, but specificity of binding impacts RNA-ligand interactions in complex, biologically relevant settings. Taken together, these results demonstrate that even though diverse ligands can bind to the same aptamer binding site, factors including selectivity, mode of recognition, and impacts on both conformational kinetics and thermodynamics can all play a role in the ability of a compound to modulate biological function by binding to RNA.

## DATA AVAILABILITY

Abasic mutant at positions 13, 14, and 15 (ab13_14_15) of the PreQ_1_ riboswitch aptamer domain from *Tte* co-crystallized with compounds **4** and **8**, have been submitted to the Protein Data Bank for the details on atomic coordinates and structure factors under the accession codes 8YAM and 8YAN respectively. The sequencing reads used for SHAPE-MaP were deposited into the NLM/NCBI Sequence Read Archive (SRA) under the BioProject ID-PRJNA1077397. All other relevant data are available from the authors upon request Source data are provided with this paper.

## ACKNOWLEDGEMENTS

The project was supported by the intramural program of the National Institutes of Health, National Cancer Institute, Center for Cancer Research (Grant 1-ZIA-BC011585-10 to J.S.S.). For MST instrumentation and data analysis, we thank Dr. Sergey G. Tarasov and Marzena Dyba from the CCR Center for Structural Biology Biophysics Resource. We also thank Professor Joseph E. Wedekind for the generous gift of the pEnv8(GAAA) plasmid that was re-engineered in the study. For the data analysis support, this project has also been funded in part with Federal funds from the NCI, National Institutes of Health, Department of Health and Human Services, under Contract No. 75N91019D00024. The content of this publication does not necessarily reflect the views or policies of the Department of Health and Human Services, nor does mention of trade names, commercial products, or organizations imply endorsement by the U.S. Government. X-ray diffraction data were collected at BL45XU of SPring-8 (Hyogo, Japan) with the approval of the Japan Synchrotron Radiation Research Institute (JASRI) under proposal number of 2021B2715. This work was supported in part by the Japan Society for the Promotion of Science (JSPS KAKENHI Grant Numbers 20K21281 and 20H02916) and a grant from The Uehara Memorial Foundation to T.N.

