## Supplemental Information for "Mechanistic Analysis of Riboswitch Ligand Interactions Provides Insights into Pharmacological Control over Gene Expression"

**Supplementary Information**

**Material and Methods:**

**Microscale Thermophoresis**

100nM *Cy5*-labeled *Bsu* PreQ_1_ (GCGGGAGAGGUUCUAGCUACACCCUCUAUAAAAAACUAAGGACGAGCUGUAUCCUUGGAUACGGCC) was prepared in 1X PreQ_1_ buffer (50 mM Tris, pH 7.5, 100 mM KCl, 1 mM MgCl_2_) and annealed by heating to 75°C for 5 min with slow cooling to room temperature (RT) for 30 mins. Serial dilutions of the compounds were prepared in 10% DMSO in 1X PreQ_1_ buffer. Equal volumes of folded RNA and compound was equilibrated for 15 min at RT. Using premium coated capillaries, MST was conducted in triplicate on a Monolith NT.115 system (NanoTemper Technologies). Obtained values were plotted against the ligand concentration and a dissociation constant (K_D_) was determined for each of the compounds using a single-site binding model.

**Fluorescence Intensity Assay**

*Cy5*-labeled *Bsu* PreQ_1_ aptamer RNA construct was purchased from IDT and annealed in PreQ_1_ folding buffer as above. Compound dilution was prepared at a concentration range of 0 to 250 μM in 1X PreQ_1_ buffer with 5% DMSO. RNA to a final concentration of 50nM is added to each well and incubated at RT for 15 min. The spectrofluorometer FluoroMax-4C ( Horiba Instruments) was used to read the fluorescence intensity at an excitation wavelength of 645 nm and an emission wavelength of 650-750 nm. The fluorescence intensities were plotted against ligand concentration. The binding affinity was calculated by fitting the curve using single-site binding model in GraphPad Prism 10.

**Single-round transcription termination assay**

The transcription termination assays were carried out as previously reported^1^. The DNA fragment containing λPR promoter, 26-nt C-less sequence followed by the *Staphylococcus saprophyticus* (*Ssa*) PreQ_1_ riboswitch, and its downstream sequence, was amplified by PCR from the plasmid. The PCR product was then gel-purified and used as the DNA template for transcription termination assay. Halted transcription complexes were prepared in a solution containing 1 µM GTP, 5 µM ATP, 5 µM UTP, 100 µM ApU, [α-^32^P] GTP, 75 nM DNA template, 0.0167 U/µL *Escherichia coli* RNA polymerase holoenzyme (New England BioLabs) in 1 × transcription buffer (20 mM Tris-HCl, pH 8.0, 2 mM NaCl, 1 mM MgCl_2_, 4% glycerol, 0.1 mM DTT, and 0.1 mM EDTA), and incubated at 37 °C for 15 min. A DNA oligonucleotide complementary to the C-less sequence was added to the reactions (final concentration of 1.1 µM) in 1 × transcription buffer and incubated at room temperature for 5 min. Subsequent elongation was restarted by combining 9 µL of halted transcription complex, 3 µL of each compound (0–5 mM compound and 25% DMSO in 1 × transcription buffer), and 3 µL of NTPs mix (200 µM each NTP, 100 µg/mL heparin, and 250 mM KCl in 1 × transcription buffer), and incubated at 37 °C for 20 min. Then, 0.5 U of RQ1 RNase-Free DNase (Promega) was added to the reactions and incubated at 37 °C for 10 min to cleave the DNA template. The reactions were stopped by adding equal volume of loading dye (8 M urea, 20% sucrose, 0.05% bromophenol blue, and 0.05% xylene cyanol in 2 × TBE). The reaction mixture was separated by 8% denaturing PAGE and visualized by phosphorimager. The band intensity was analyzed by ImageQuant software (GE Healthcare). Termination efficiency was calculated with dividing the intensity for the terminated RNA band by those for the total (terminated and antiterminated) RNAs.

**X-ray co-crystal structure determination and refinement**

Abasic mutant at positions 13, 14, and 15 (ab13_14_15) of the PreQ_1_ riboswitch aptamer domain from *Thermoanaerobacter tengcongensis* (*Tte*) was co-crystallized either with the compound **4** or **8**, under the conditions containing 100–300 mM potassium sodium tartrate, 100 mM sodium citrate (pH 5.6), and 2.0–2.6 M ammonium sulfate, at 20 °C. The crystals were soaked into the cryoprotectant solution containing 60 mM potassium sodium tartrate, 25 mM sodium citrate (pH 5.6), and 2.15 M lithium sulfate and then flash frozen by plunging into liquid nitrogen. X-ray diffraction data were collected at the beamline BL45XU of the SPring-8 (Hyogo, Japan) with the aid of an automatic data-collection system ZOO^2^. Diffraction data were integrated and scaled with the programs KAMO^3^ and XDS^4^. Data processing statistics are summarized in Table S4. The structures were determined by the molecular replacement method with the program PHASER^5^, using the structure of *Tte* PreQ_1_ aptamer domain (PDB ID: 6E1S) as a search model. The solutions were first subjected to rigid body, simulated annealing, energy minimization, restrained isotropic B-factor refinement with PHENIX^6^, generating clear electron density maps corresponding to each compound. The atomic models were built manually with COOT^7^ and improved by iterative cycles of refinement with PHENIX. The current co-crystal structures with the compounds **4** and **8** were refined to an *R*_free_ of 0.193 and 0.214 at 2.15 and 2.25 Å resolution, respectively. Atomic coordinates and the structure factors of the co-crystal structures with the compounds **4** and **8** have been deposited in the Protein Data Bank, under the accession codes 8YAM and 8YAN, respectively.

**Structure probing of different riboswitches using SHAPE-MaP**

*In vitro* transcription for RNA preparation: Ultramer® DNA Oligos purchased from IDT (Table S1) were amplified using Q5® High-Fidelity DNA Polymerase (NEB) following manufacturer’s instructions and used as templates for *in vitro* transcription. A list of primers used for template amplification can be found in Table S2. Using the HiScribe™ T7 High Yield RNA Synthesis Kit (NEB), RNA was transcribed following the manufacturer’s guidelines and purified using Zymo RNA Clean and Concentrator−5 kit (Zymo).

Acylation of RNA *in vitro* using SHAPE reagent: 10pM of the purified RNA in 12μL of nuclease-free water was denatured by heating at 95 °C for 2 min and snap-cooled on ice. 3.3X PreQ_1_ folding buffer (165mM Tris, pH 7.5, 333mM KCl, 3.3mM MgCl2) was used for folding the RNA by incubating at 75 °C for 5 min and slowly cooled to RT for 30 min. The folded RNA was incubated with DMSO (control), PreQ_1_ (cognate ligand) and **4** (synthetic ligand) with a final concentration of 2.5%, 10 μM and 150 μM respectively, for another 15 min.^8^ For acylation, 5 pmol of each RNA was either treated with 1μl DMSO (-RNA) or was chemically modified using 100mM 2A3 (+RNA) by incubation at 37 °C for 20 min. The acylation was quenched using 1M DTT and purified using Illustra G-25 columns.

Reverse transcription of RNA for Mutational profiling (MaP): 1μl of dNTPs (10mM each, NEB) and 1 μl RT oligo (20 μM) (Table S2) were added to +/- RNA followed by incubation at 70°C for 5 min and snap cooled for a min. Next, first strand buffer mix containing 4 μl of (5X, 250mM Tris-HCl pH 8.0, 375mM KCl), 2 μl DTT (100mM), 1 ul RNase Inhibitor, 1 ul SuperScript II (Invitrogen), 1 ul MnCl_2_ (120 mM) was added, followed by incubation at 25°C for 5 min. The +/1 samples were reverse transcribed by incubating at 42°C for 2 hours followed by heat-inactivation of the enzyme by incubating at 75°C for 20 min. The resulting cDNA were purified using illustra G-25 spin columns.

Library generation and data analysis: The libraries were generated by the PCR amplification (Q5® High-Fidelity DNA Polymerase, NEB) of the above cDNA using two sets of primer pairs (Table S2). The libraries were purified using DNA clean and concentrator-5 (Zymo) 1pM of each library was pooled for paired-end sequencing on an Illumina Miniseq instrument following manufacturer’s instructions. SHAPEmapper software ([https://github.com/Weeks-UNC/shapemapper2/tree/master](about:blank)) using default parameters was used to generate SHAPE profiles for each of the RNA. Delta SHAPE analysis ([https://github.com/Weeks-UNC/deltaSHAPE](about:blank)) was performed on the generated .map files. RNA structure prediction using SHAPE constraints was done on structure editor ([https://rna.urmc.rochester.edu/GUI/html/StructureEditor.html](about:blank)). Superfold was used to generate the arc plots ([https://github.com/Weeks-UNC/Superfold](about:blank)).

***In vivo* GFPuv reporter assay**

The construct for *Ssa-*PreQ_1_ riboswitch-GFPuv reporter assay was designed following Dutta et al, 2018.^9^ Successful insertion of the desired RNA sequence was confirmed using Sanger sequencing. The construct was then transformed into competent *E. coli* strain JW2765 *ΔqueF* (Coli Genetic Stock Center, Yale University). Cells were streaked in the presence and absence of DMSO (control), PreQ_1_(natural ligand), **4**, **8**, **9** (synthetic ligands), and empty vector and were grown on specialized CSB agar media at 37 °C overnight. The plates were visualized for GFP fluorescence on a UV-transilluminator at 365nm and photographed.

**Table S1: List of riboswitch constructs**

|  | *T7 promoter*-**aptamer sequence**-RT primer binding site |
| --- | --- |
| *B. subtilis* | *TAATACGACTCACTATAG*GGCCTTCGGGCCAAGCGGG**AGAGGTTCTAGCTACACCCTCTATAAAAAACTAA**GGACGAGCTGTATCCTTGGATACGGCCTTTTTCGATCCGGTTCGCCGGATCCAAATcgggcttcggtccggttc |
| *T. tencongensis* | *TAATACGACTCACTATAG*GGCCTTCGGGCCAACTCAC**CTGGGTCGCAGTAACCCCAGTTAACAAAACA**AGGGAGGTAATTTTTCGATCCGGTTCGCCGGATCCAAATcgggcttcggtccggttc |
| *S. saprophyticus* | *TAATACGACTCACTATAG*GGCCTTCGGGCCAATATAA**AGAGGTTCCTAGCTGATACCCTCTATAAAAAACTA**GACACATGTACAACGTCTGTCTTTTTTATAGAGATAGGCGTTTTTTCGATCCGGTTCGCCGGATCCAAATcgggcttcggtccggttc |
| *L. rhamnosus* | *TAATACGACTCACTATAG*GGCCTTCGGGCCAAGGAACCGCCCCGCGCGCTTCCACGATACTTATTTCCTTTGATCGTCGTTATTACTGGCAAAGCCACAAAGGAGTCGATCCGGTTCGCCGGATCCAAATcgggcttcggtccggttc |
| *F. prausnitzii* | *TAATACGACTCACTATAG*GGCCTTCGGGCCAAGGAACGCTGAAAGAGCAACTTAGGATTTTAGGCTCCCCGGCGTGTCTCGAACCATGCCGGGCCAAACCCATAGGGCTGGCGGTCCCTGTGCGGTCAAAATTCATCCGCCGGAGTATTGTTATCGATCCGGTTCGCCGGATCCAAATcgggcttcggtccggttc |
| *S.  pneumoniae* | *TAATACGACTCACTATAG*GGCCTTCGGGCCAACTTGGTGCTTAGCTTCTTTCACCAAGCATATTACACGCGGATAACCGCCAAAGGAGAATCGATCCGGTTCGCCGGATCCAAATcgggcttcggtccggttc |

**Table S2: Primer sequences**

| *In vitro* transcription  SHAPE_temp_F  SHAPE_temp_R | TAATACGACTCACTATAGGGCC  GAACCGGACCGAAGCC |
| --- | --- |
| SHAPE PCR1:  Forward primers: SHAPE_PCR1_*Bsu*:  SHAPE_PCR1_*Tte*:  SHAPE_PCR1_*Ssa*:  SHAPE_PCR1_*Lrh*:  SHAPE_PCR1_*Fpn*:  SHAPE_PCR1_*Spn*:  Reverse primer : | GACTGGAGTTCAGACGTGTGCTCTTCCGATCTNNNNNGCGGGAGAGGTTCTAGCTAC  GACTGGAGTTCAGACGTGTGCTCTTCCGATCTNNNNNCTCACCTGGGTCGCAGTAAC  GACTGGAGTTCAGACGTGTGCTCTTCCGATCTNNNNNTATAAAGAGGTTCCTAGCTG  GACTGGAGTTCAGACGTGTGCTCTTCCGATCTNNNNNGGAACCGCCCCGCGCGCTTC  GACTGGAGTTCAGACGTGTGCTCTTCCGATCTNNNNNGGAACGCTGAAAGAGCAACT  GACTGGAGTTCAGACGTGTGCTCTTCCGATCTNNNNNCTTGGTGCTTAGCTTCTTTC  CCCTACACGACGCTCTTCCGATCTNNNNNGAACCGGACCGAAGCCCG |

**RNA Sequences and Preparation for MAGNA analysis**

The *Bsu* PreQ_1_ sequence (34nt) was flanked by 6 nucleotides of single-stranded RNA and a sequence that is used to allow the annealing to the DNA handles for surface (3’ DNA splint in Table S3) and bead (5’ DNA splint in Table S3) attachment. The RNA was synthesized by IDT and tagged with 5’ biotin. The DNA handles were synthesized by Eurofins.

The RNA was annealed to the two DNA handles (10 µM) in Annealing Buffer (10mM Tris-HCl, pH7.4, 50 mM NaCl, 1mM EDTA). Once annealed, the structures were mixed with an equal volume of GenTegraRNA (GenTegra) and were stored at -20°C until use. The annealed RNA structure (2 fmol) was then hybridized to 3 µl of MyOne T1 streptavidin beads (Invitrogen) in Hybridization Buffer (10% PEG8000, 5xSSC) for 10 min before being resuspended in Oligo Buffer (OB) (1xPBS, 0.2% BSA, 0.1% NaAz) for loading in the instrument.

**Table S3:**

| *Bsu* PreQ_1_ RNA | 5’biotin-UCGGCGAUCUACGCAGCGACAUAAUAAGAGGUUCUAGCUACACCCUCUAUAAAAAACUAAUAACAACCACUUCCUAAUCUGUCAUCUUCUG |
| --- | --- |
| 5’ DNA splint | GTCGCTGCGTAGATCGCCGA |
| 3’ DNA splint | GTGTCTTTTGGTCTTTCTGGTGCTCTTCGAATCAGAAGATGACAGATTAGGAAGTGG |

**Instrument and Measurement**

Flow cell surfaces were functionalized with an oligonucleotide and then passivated with OB buffer after flow cell assembly. The RNA bound to the MyOne beads were injected into the flow cell and allowed to hybridize to the surface bound oligonucleotide for 30 minutes in OB. After hybridization, the buffer was changed to the testing buffer (50 mM HEPES, pH7.4; 100 mM KCl; 1mM MgCl_2_; 1% DMSO). Unbound beads were removed by washing and any non-specifically bound beads were removed by increasing the magnetic force to ~25-30 pN. Subsequent experiments were performed at 22°C, with recording of bead vertical or Z-positions at 30Hz.

Magnetic force spectroscopy was performed using Depixus’ prototype MAGNA instrument. ^10^ A force ramp experiment (approximately 1-25 pN) was first recorded for each structure in test buffer containing 1% DMSO. This experiment was used as a control for normalization as well as to identify structures that could be opened and closed. Next **4** or PreQ_1_ ligand was added at increasing concentration in testing buffer. For each condition, 50 ramp cycles were recorded. Stepped constant-force experiments were also performed. To do so, the applied magnetic force was held constant for a fixed time (~30 sec) before increasing it stepwise. For constant-force experiments, the beads were submitted to the same force for 30-60min.

**Analysis of ramp experiment data**

We only analyzed the signal of beads whose measured variance in position was roughly equal to the noise expected from Brownian motion. Signals with excessive tracking noise or detached beads were thus excluded. For each structure, each cycle was analyzed separately and the size and corresponding applied force of abrupt jumps in bead Z-position due to unfolding and refolding of the RNA structure were detected using an algorithm based on clustering the force-extension data with the HDBScan python module. ^11^ We first selected parameters to define analyzable RNA structures: 1) An unfolding force between 5-15 pN and an unfolding size between 10-25 nm in control (DMSO) conditions; 2)Total number of analyzable cycles >20; 3) The proportion of cycles with an unfolding event >80%; and 4). The structures were present in more than half of the conditions tested. Structures were considered folded when analyzable cycles contained no unfolding or refolding events.

For each analyzable structure, the median forces at which unfolding and refolding occurred in DMSO conditions were used to normalize all cycles of all other conditions. The normalized unfolding and refolding forces were plotted to study the changes in distribution with different concentrations of **4** and the median of the force at the maximum peak of the distribution for each structure was plotted against concentration of **4**. For PreQ_1_ ligand binding, a threshold force corresponding to the 0.95 percentile of the unfolding force distribution was determined using DMSO conditions. The fraction of force ramp cycles with an unfolding force over the threshold for each condition or did not unfold was calculated and plotted against the PreQ_1_ ligand concentration. For both **4** and PreQ_1_ ligand, K_D_/EC_50_ was calculated using the following formula: y=(u+b*x/K_D_)/(1+x/K_D_).

**Analysis of constant-force experiments**

Data were excluded from RNA structures showing high noise (i.e., a variation in extension exceeding twice the expected length of the structure within a 1*s* window), or structures that did not fold and unfold in response to the applied magnetic force or that did not form stable folded structures in the presence of PreQ_1_ ligand. A simple heuristic Hidden-Markov-Model was then used to attribute a state (folded or unfolded) to each data point (the frequency of data collection was 30 Hz) with the data being presented as a trace of bead position versus time. This allowed the time the structure spent consecutively in either state before re-/unfolding (lifetime) to be assessed. The distribution of lifetimes for each individual structure and state were then analyzed and Maximum Likelihood Estimation (MLE) was used to estimate whether one, two or three exponential distributions were present. The Bayesian information criterion (BIC) was used to confirm that for all conditions, a single exponential distribution was the preferred model, except for the folded state lifetimes when the PreQ_1_ ligand was present for which a mixture of two exponential distributions with two distinct lifetimes was the preferred model. To assess ligand concentration dependency, we computed the mean observed lifetime of the folded and unfolded states for the DMSO control and different concentrations of **4**. For the PreQ_1_ ligand, we used the lifetimes obtained from the MLE analysis , together with the weights of the two distributions. To aggregate data from multiple beads, it was necessary to first compensate for minor differences in paramagnetic bead magnetism by normalizing each RNA structure with its control condition. This was achieved through analyzing the logarithm of un-/refolding rates log(1/lifetime) which according to the Kramers-Bell theory^12^ scales linearly with the applied force.

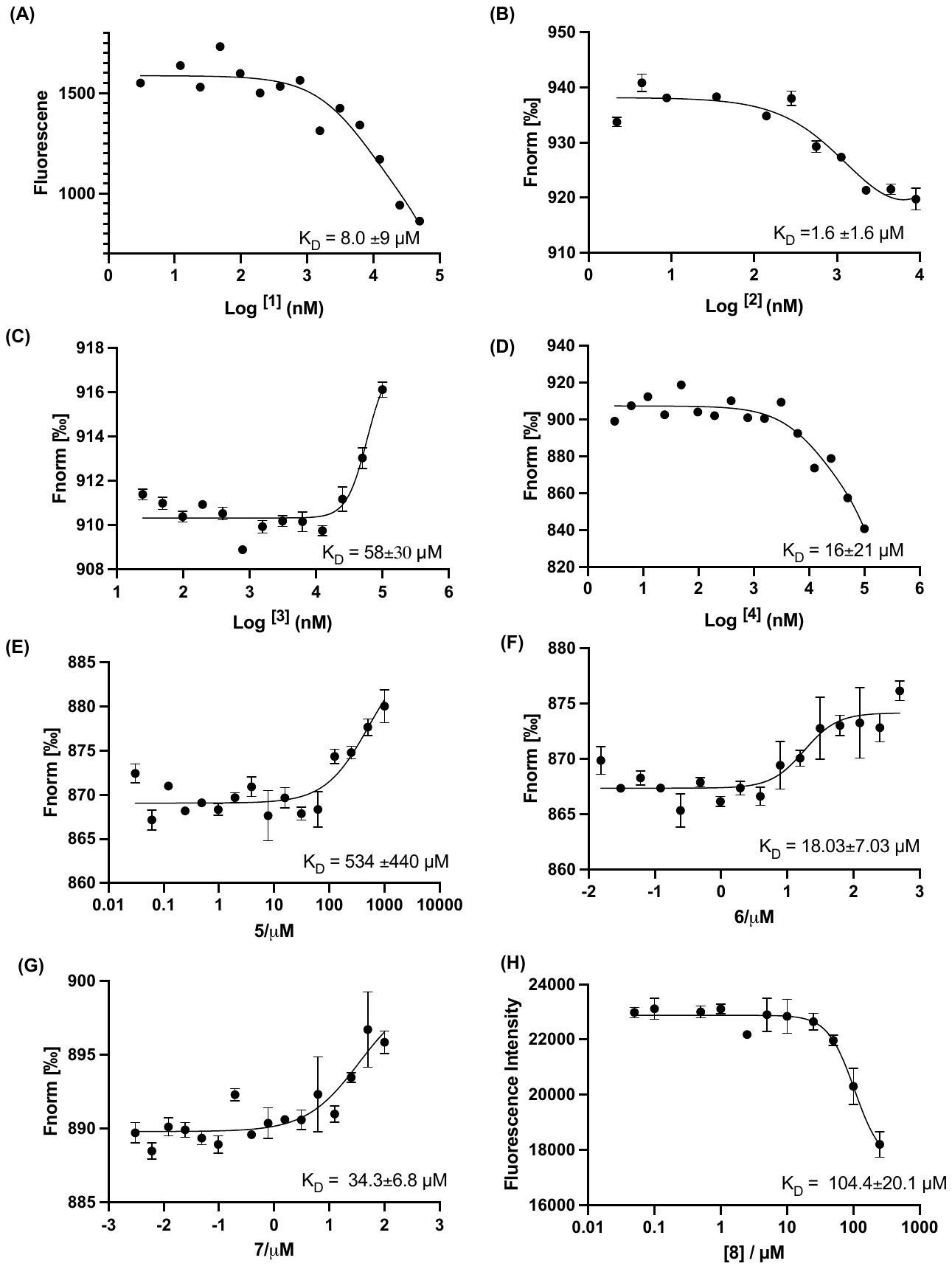

**Figure S1.**  Affinity measurements of PreQ_1_ ligands derivatives to *Bsu*-PreQ_1_ riboswitch. Compounds **1 (**A), **2** (B), and **4** (D) show inherent fluorescence titration with increasing concentrations. MST of Cy5-labeled RNA with increasing concentrations of compounds **3** (C), **5** (E) ,**6** (F) ,**7** (G) and fluorescence intensity with increasing concentrations of **8** (H). Error bars indicate the standard deviation determined from three independent measurements.

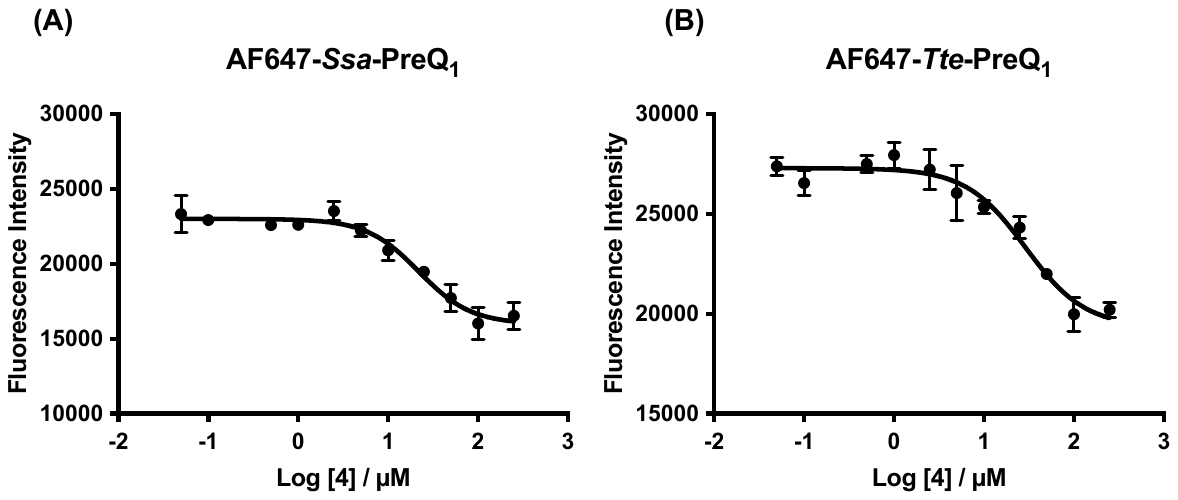

**Figure S2.** Affinity measurement using fluorescence titrations of 5′-AlexaFluor 647 labeled (A) *Ssa-*PreQ_1_ RNA and (B) *Tte-*PreQ_1_ RNA aptamers in the presence of increasing concentration of **4**. Error bars indicate the standard deviation determined from three independent measurements.

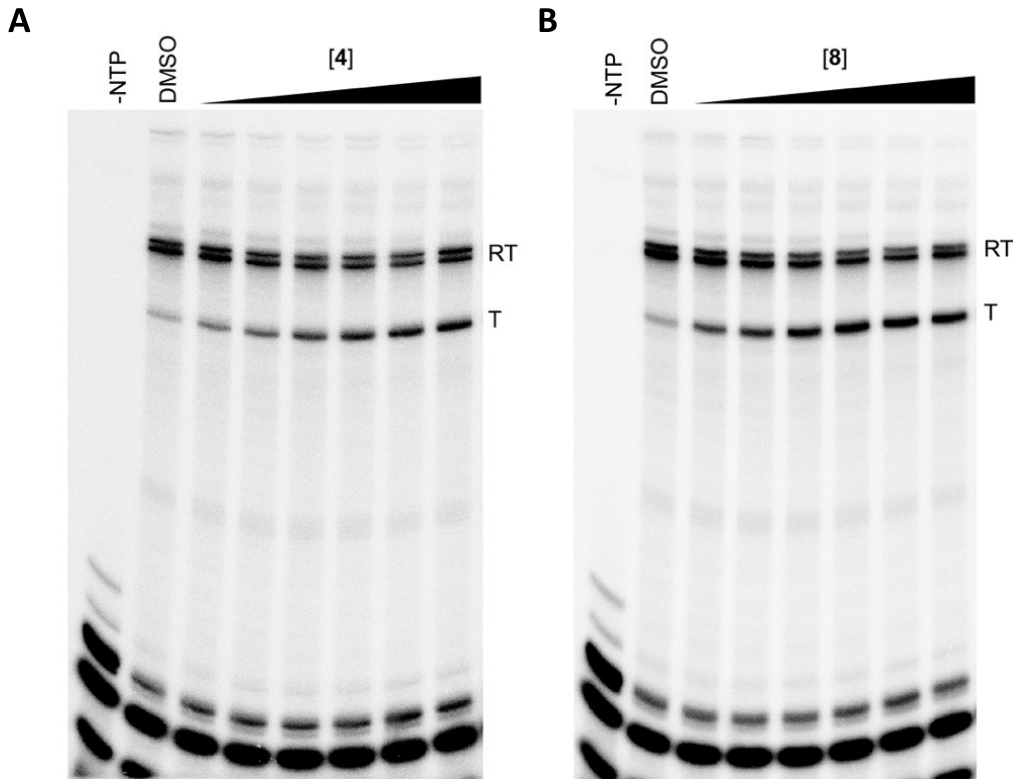

**Figure S3.** Transcription termination assay on *Ssa*-PreQ_1_ riboswitch in the presence of increasing concentrations of **4** (A) and **8** (B) demonstrating functional activity *in vitro*. RT: Full transcript read-through, T: transcription halt due to termination. Shown are the representative PAGE gels from three independent replicates and transcriptional termination efficiency (T_50_) value is quantified on the right using [Agonist] vs response – variable slope (four parameters) model. Error bars on T_50_ values represent standard deviation from the replicates.

**Table S4**. Summary of data collection and refinement statistics.

|  | ab13_14_15-**4** | ab13_14_15-**8** |
| --- | --- | --- |
| **Data collection** |  |  |
| Space group | *P*6_3_22 | *P*6_3_22 |
| Cell dimensions |  |  |
| *a, b, c* (Å) | 115.9, 115.9, 58.3 | 116.4, 116.4, 57.8 |
| Wavelength (Å) | 1.0 | 1.0 |
| Resolution (Å) | 41.1-2.15 (2.28-2.15) | 41.0-2.25 (2.39-2.25) |
| *R*_merge_^a^ | 0.083 (3.36) | 0.064 (3.62) |
| *R*_p.i.m._^b^ | 0.015 (0.585) | 0.012 (0.641) |
| CC_1/2_^c^ | 0.999 (0.595) | 0.999 (0.589) |
| <*I*>*/*<s*I*> | 29.8 (1.57) | 32.4 (1.44) |
| Completeness (%) | 99.6 (99.2) | 99.9 (99.9) |
| Redundancy | 33.2 (33.3) | 31.2 (31.9) |
| **Refinement** |  |  |
| Resolution (Å) | 41.1-2.15 | 41.0-2.25 |
| No. reflections | 12,983 | 11,405 |
| *R*_work_^d^/*R*_free_^e^ | 0.184/0.193 | 0.198/0.214 |
| No. atoms |  |  |
| RNA | 667 | 663 |
| Ligand | 21 | 20 |
| Water | 13 | 5 |
| *B*-factors (Å^2^) |  |  |
| RNA | 77.5 | 97.6 |
| Ligand | 68.6 | 75.0 |
| Water | 70.2 | 87.4 |
| R.m.s. deviations |  |  |
| Bond lengths (Å) | 0.005 | 0.002 |
| Bond angles (º) | 0.982 | 0.372 |
| PDB ID | 8YAM | 8YAN |

The values in parentheses are for the outermost shell.

^a^ *R*_merge_ = S*_hkl_* S*_i_*|*I_i_*(*hkl*) – <*I*(*hkl*)>|/S*_hkl_* S*_i_* *I*_i_(*hkl*), where *I*_i_(*hkl*) is the observed intensity and S*I*(hkl)> is the average intensity over symmetry-equivalent measurements.

^b^ Definition of *R*_p.i.m._^13^

^c^ Pearson correlation coefficient between intensities of random half-dataset. ^14^

^d^ *R*_work_ = S|*F*_o_ – *F*_c_|/S*F*_o_ for reflections of working set.

^e^ *R*_free_ = S|*F*_o_ – *F*_c_|/S*F*_o_ for reflections of test set (5.0% of total reflections).

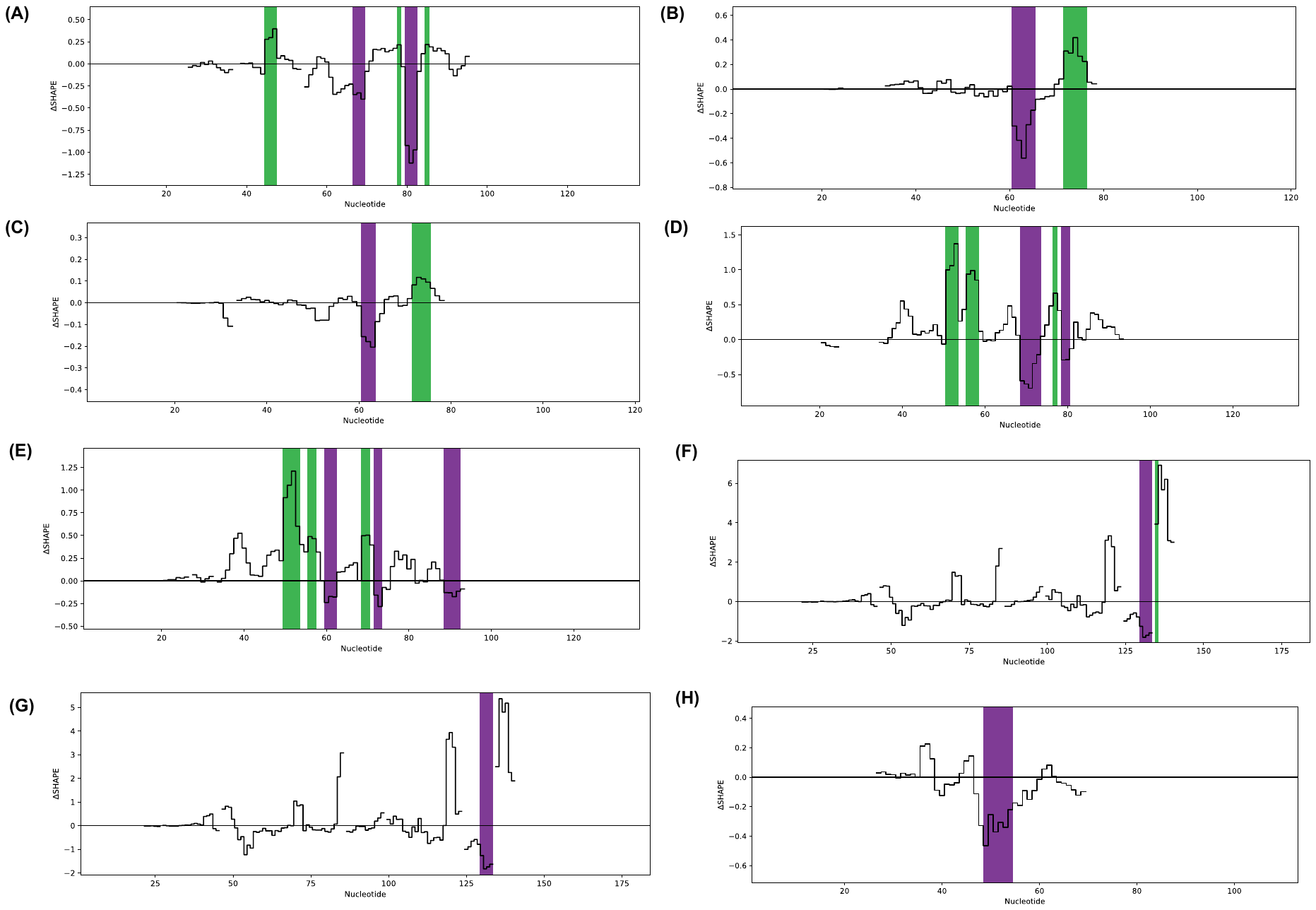

**Figure S4.** Delta SHAPE-MaP profiles of (A) *Bsu-*PreQ_1_ in presence of PreQ_1_ ligand, *Spn-*PreQ_1_ with PreQ_1_ (B) and **4** (C), *Lrh-*PreQ_1_ with PreQ_1_ (D) and **4** (E), *Fpn*-PreQ_1_ with PreQ_1_ (F) and **4** (G) and *Tte-*PreQ_1_ in the presence of PreQ_1_ ligand (H). Shown here are the delta SHAPE profiles with significant changes only.

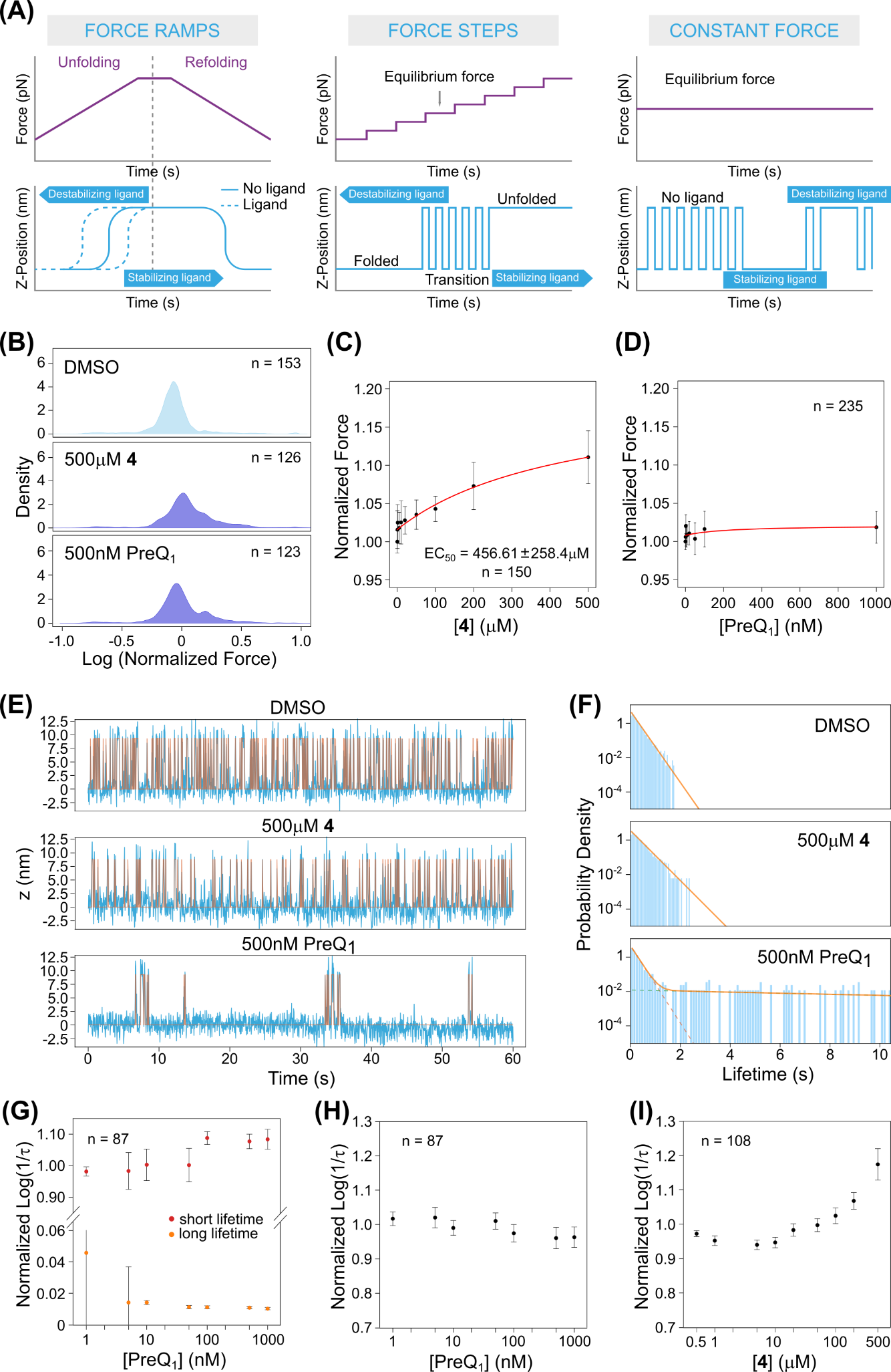

**Figure S5.** (A) Schema showing the different MAGNA experimental modes used. In ramp experiments the molecules are subjected to cycles of gradually increased and decreased force, whilst in stepped constant force experiments the force is held constant for a fixed period before being increased stepwise. In constant force experiments the molecules are subjected to a single constant force for an extended period, allowing the transition between different states to be observed. The effect of the binding of stabilizing and destabilizing ligands are indicated. (B) The normalized force distribution for the refolding force, comparing DMSO (control), 500 µM **4** and 500 nM PreQ_1_ ligand. (C-D) Dose-response curve of the refolding force with increasing concentration of **4** (C) and PreQ_1_ (D). (E) Another example of the raw trace of a constant force experiment for a single molecule in the presence of DMSO, 500 µM **4** and 500 nM PreQ_1_. The force is held at the equilibrium force, and the Z-position is plotted again the time. (F) Fitting of the probability density of folded lifetimes of a single molecule in constant force experiments. The probability in log scale was plotted again time, and the overall fitting is shown as orange lines. In the PreQ_1_ condition, two distributions can be fitted, depicted as the two dotted lines. (G) The normalized unfolding rate (1/ folded lifetime) with increasing concentration of PreQ_1_ ligand. Both the short- and the long-folded lifetimes are plotted in the same graph. (H-I) The refolding rate (1/unfolded lifetime) of the PreQ_1_ riboswitch with increasing concentrations of PreQ_1_ (H) or **4** (I), from constant force experiments.

**General chemistry methods:**

Unless otherwise noted, all chemical reagents were obtained from commercial suppliers and used without further purification. Solvents were removed using a Buchi rotary evaporator under reduced pressure. Flash column chromatography was performed using a Teledyne ISCO CombiFlash Rf automated chromatography system.

High resolution mass spectrometry data were acquired on an Agilent 6520 Accurate-Mass Q-TOF LC/MS System, (Agilent Technologies, Inc.) equipped with a dual electro-spray source, operated in the positive-ion mode. Separation was performed on Zorbax 300SB-C18 Poroshell column (2.1 mm x 150 mm; particle size 5 mm). The analytes were eluted using a water/acetonitrile gradient with 0.1% formic acid.  Data were acquired at high resolution (1,700 *m/z*), 4 GHz. To maintain mass accuracy during the run time, an internal mass calibration sample was infused continuously during the LC/MS runs. Data acquisition and analysis were performed using MassHunter Workstation Data Software, LCMS Data Acquisition (version B.06.01) and Qualitative Analysis (version B.07.00).

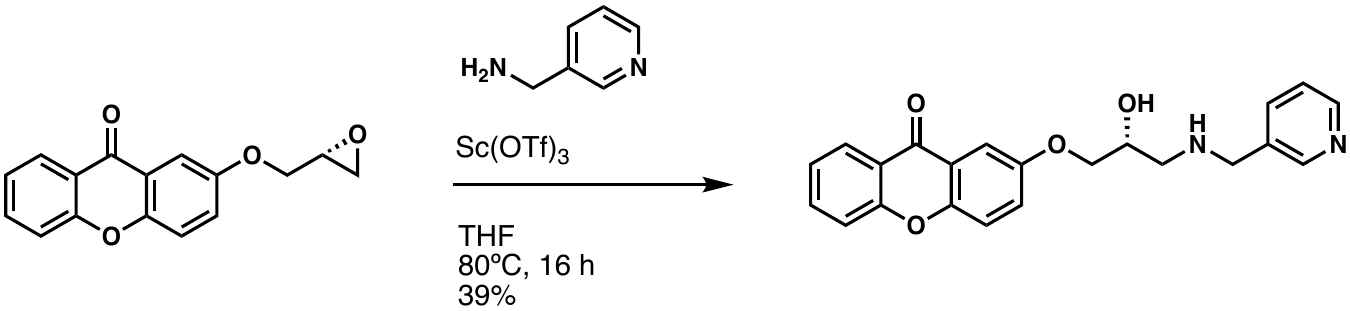

To a solution of 2-(oxiran-2-ylmethoxy)xanthen-9-one (40 mg, 0.15 mmol) in THF (3 mL) was added 3-pyridinemethanamine (0.018 mL) and scandium (III) triflate (4 mg). The mixture was heated to 80ºC and stirred for 16 hours, then was returned to room temperature. The mixture was concentrated down in vacuo, and the crude product was purified by HPLC to afford 22 mg of the desired product (39%). ^1^H NMR (500 MHz, DMSO-*d*_6_): δ 9.20 (d, *J* = 30.9 Hz, 2H), 8.72 (d, *J* = 49.8 Hz, 2H), 8.20 (dd, *J* = 7.9 Hz, 1H), 8.05 (d, *J* = 7.8 Hz, 1H), 7.90-7.86 (m, 1H), 7.68 (d, *J* = 9.1 Hz, 1H), 7.67 (d, *J* = 8.2 Hz, 1H), 7.60 (d, *J* = 3.2 Hz, 1H), 7.56 (q, *J* = 4.8, 3.0 Hz, 1H), 7.51-7.47 (m, 2H), 4.31 (s, 2H), 4.28-4.23 (m, 1H), 4.14-4.09 (m, 2H), 3.23 (s, 1H), 3.08 (s, 1H); ^13^C NMR (125 MHz, DMSO-*d*_6_): δ 175.8, 155.6, 154.7, 150.6, 150.5, 149.4, 146.8, 139.0, 135.5, 128.2, 126.0, 125.0, 124.3, 121.5, 120.5, 119.9, 118.2, 107.0, 70.4, 64.9, 49.0, 47.8; HRMS: (ESI+) m/z calculated for C_22_H_20_N_2_O_4_ [M+H]^+^: 377.1496, found: 377.1496.

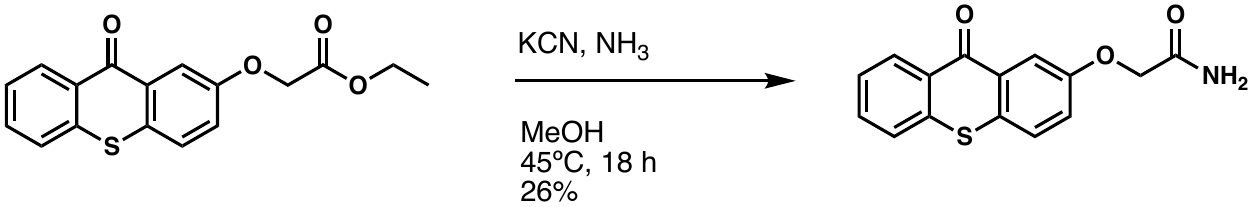

In a sealed microwave vial, a solution of ethyl 2-[(9-oxo-9H-thioxanthen-2-yl)oxy]acetate (50 mg, 0.16 mmol), potassium cyanide (2 mg, 0.032 mmol), and ammonia in methanol (2 mL) was heated to 45ºC and stirred for 18 hours. The mixture was returned to room temperature, then the vial and solution were purged with nitrogen. After removing the solvent in vacuo, the resulting crude product was redissolved in dichloromethane and washed with water (10 mL). The organics were extracted with dichloromethane (3 x 10 mL), washed with brine (10 mL), then dried over MgSO_4_. The crude product was concentrated down onto silica gel, then purified by ISCO Flash column chromatography (DCM:MeOH, 0-40%) to afford 12 mg of the desired product (26%). ^1^H NMR (500 MHz, DMSO-*d*_6_): δ 8.48 (dd, *J* = 8.2 Hz, 1H), 7.93 (d, *J* = 2.9 Hz, 1H), 7.86 (dd, *J* = 8.1 Hz, 1H), 7.84 (d, *J* = 8.8 Hz, 1H), 7.79-7.76 (m, 1H), 7.67 (s, 1H), 7.61-7.57 (m, 1H), 7.49 (dd, *J* = 8.8 Hz, 1H), 7.43 (s, 1H), 4.60 (s, 2H); ^13^C NMR (125 MHz, DMSO-*d*_6_): δ 178.4, 169.4, 156.7, 136.8, 132.8, 129.4, 129.1, 128.8, 128.1, 127.7, 126.7, 126.6, 122.8, 111.7, 66.8; HRMS: (ESI+) m/z calculated for C_15_H_11_NO_3_S [M+H]^+^: 286.0532, found: 286.0531.

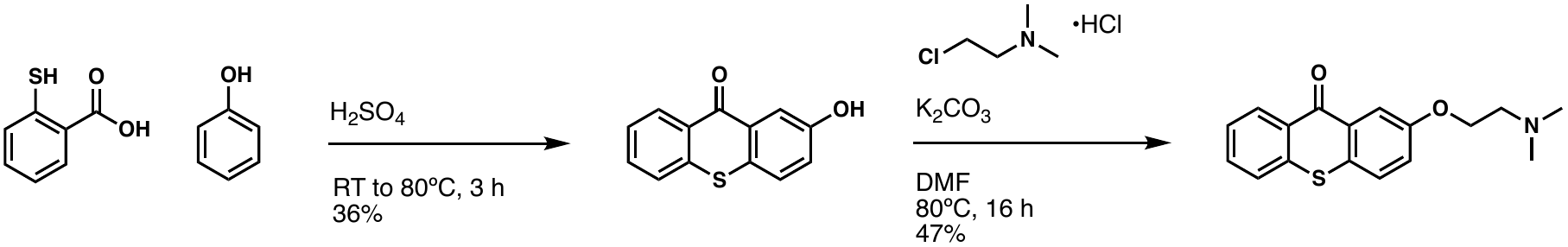

2-Mercaptobenzoic acid (4 g, 25.9 mmol) was slowly added to sulfuric acid (40 mL) at room temperature in a round bottom flask equipped with a magnetic stir bar. The mixture stirred for 15 minutes, then phenol (9.8 mL, 111.5 mmol) was slowly added over 15 minutes. The reaction mixture was then heated to 80ºC and continued stirring for 3 hours. The reaction was diluted with 700 mL of boiling water then allowed to cool to room temperature. The resulting precipitate was then collected by filtration to obtain 2.14 g of 2-hydroxythioxanthone (36%). ^1^H NMR (500 MHz, DMSO-*d*_6_): δ 8.46 (dd, *J* = 7.9 Hz, 1H), 7.86 (d, *J* = 2.8 Hz, 1H), 7.82 (dd, *J* = 8.0 Hz, 1H), 7.77-7.73 (m, 1H), 7.70 (d, *J* = 8.7 Hz, 1H), 7.58-7.54 (m, 1H), 7.28 (dd, *J* = 8.0 Hz, 1H); ^13^C NMR (125 MHz, DMSO-*d*_6_): δ 178.6, 137.0, 132.6, 129.6, 129.1, 128.1, 128.0, 127.8, 126.5, 126.4, 126.0, 122.8, 113.2; HRMS: (ESI+) m/z calculated for C_13_H_8_O_2_S [M+H]^+^: 229.0318, found: 229.0319.

To a solution of 2-hydroxythioxanthone (30 mg, 0.131 mmol) in DMF (3 mL) was added 2-chloro-N,N-dimethyl-ethanamine hydrochloride (19 mg, 0.131 mmol) and potassium carbonate (40 mg, 0.288 mmol). The mixture was heated to 80ºC and stirred for 16 hours. After cooling to room temperature, the reaction was diluted with water (10 mL), and the mixture was extracted with dichloromethane (3 x 10 mL). The combined organic layer was then washed with water (10 mL) and brine (10 mL), then dried with MgSO_4_. The crude product was concentrated down onto silica gel and purified by ISCO Flash Column Chromatography (DCM:MeOH, 5-40%) to afford 18 mg of the desired product (47%). ^1^H NMR (500 MHz, DMSO-*d*_6_): δ 8.48 (dd, *J* = 8.2 Hz, 1H), 7.94 (d, *J* = 2.9 Hz, 1H), 7.85 (dd, *J* = 8.1 Hz, 1H), 7.81-7.75 (m, 2H), 7.61-7.57 (m, 1H), 7.45 (dd, *J* = 8.7 Hz, 1H), 4.20 (t, *J* = 5.7 Hz, 2H), 2.68 (t, *J* = 5.7 Hz, 2H), 2.24 (s, 6H); ^13^C NMR (125 MHz, DMSO-*d*_6_): δ 178.5, 157.4, 143.4, 136.8, 132.8, 129.5, 129.1, 128.2, 127.8, 126.6, 126.6, 122.8, 111.1, 66.3, 57.5, 45.6; HRMS: (ESI+) m/z calculated for C_17_H_17_NO_2_S [M+H]^+^: 300.1053, found: 300.1058.

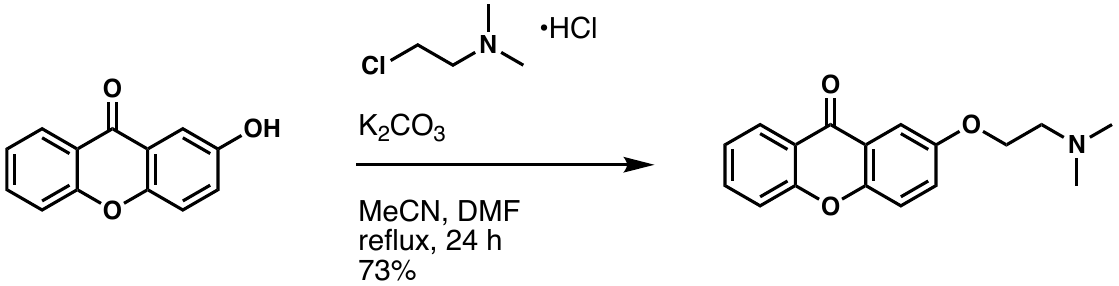

To a solution of 2-hydroxyxanthone (33 mg, 0.156 mmol) in 1:1 acetonitrile and DMF (5 mL) was added 2-chloro-N,N-dimethyl-ethanamine hydrochloride (27 mg, 0.187 mmol) and potassium carbonate (26 mg, 0.187 mmol). The mixture was heated to reflux and stirred for 24 hours. Once cooled to room temperature, the reaction mixture was diluted with water (10 mL), and the mixture was extracted with dichloromethane (3 x 10 mL). The combined organic layer was then washed with water (10 mL) and brine (10 mL), then dried with MgSO_4_. The crude product was concentrated down onto silica gel and purified by ISCO Flash Column Chromatography (DCM:MeOH, 5-40%) to afford 32 mg of the desired product (73%). ^1^H NMR (500 MHz, DMSO-*d*_6_): δ 8.20 (dd, *J* = 7.9 Hz, 1H), 7.89-7.85 (m, 1H), 7.65 (t, *J* = 9.1 Hz, 2H), 7.57 (d, *J* = 3.2 Hz, 1H), 7.50-7.45 (m, 2H), 4.16 (t, *J* = 5.7 Hz, 2H), 2.67 (t, *J* = 5.7 Hz, 2H), 2.23 (s, 6H); ^13^C NMR (125 MHz, DMSO-*d*_6_): δ 175.7, 155.5, 154.9, 150.2, 135.3, 125.9, 124.9, 124.2, 121.5, 120.5, 119.7, 118.1, 106.5, 66.4, 57.5, 45.5; HRMS: (ESI+) m/z calculated for C_17_H_17_NO_3_ [M+H]^+^: 284.1281, found: 284.1282.

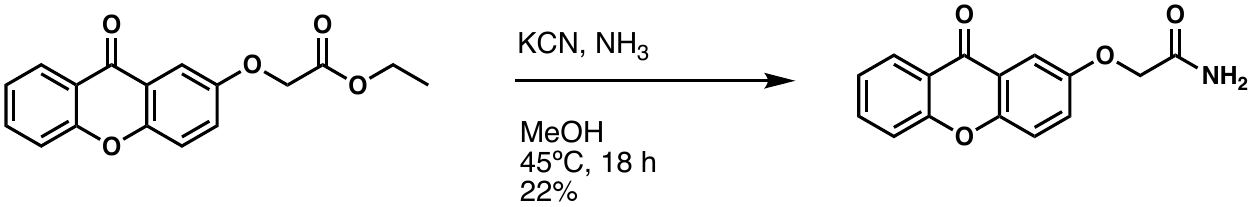

In a sealed microwave vial, a solution of ethyl 2-[(9-oxo-9H-xanthen-2-yl)oxy]acetate (50 mg, 0.17 mmol), potassium cyanide (2.2 mg, 0.034 mmol), and ammonia in methanol (2 mL) was heated to 45ºC and stirred for 18 hours. The mixture was returned to room temperature, then the vial and solution were purged with nitrogen. After removing the solvent in vacuo, the resulting crude product was redissolved in dichloromethane and washed with water (10 mL). The organics were extracted with dichloromethane (3 x 10 mL), washed with brine (10 mL), then dried over MgSO_4_. The crude product was concentrated down onto silica gel, then purified by ISCO Flash column chromatography (DCM:MeOH, 0-40%) to afford 10 mg of the desired product (22%). ^1^H NMR (500 MHz, DMSO-*d*_6_): δ 8.21 (dd, *J* = 8.1 Hz, 1H), 7.90-7.87 (m, 1H), 7.70-7.66 (m, 3H), 7.58-7.54 (m, 2H), 7.50-7.47 (m, 1H), 7.42 (s, 1H), 4.58 (s, 2H); ^13^C NMR (125 MHz, DMSO-*d*_6_): δ 175.8, 169.5, 155.6, 154.3, 150.5, 135.5, 126.0, 125.0, 124.3, 121.4, 120.5, 119.8, 118.2, 107.3, 67.1; HRMS: (ESI+) m/z calculated for C_15_H_11_NO_4_ [M+H]^+^: 270.0761, found: 270.0757.

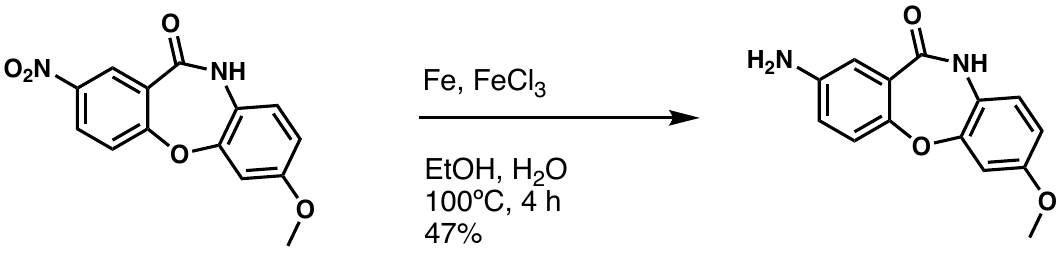

In a round bottom flask, a solution of 7-methoxy-2-nitrodibenz[b,f][1,4]oxazepin-11(10H)-one (150 mg, 0.52 mmol), iron (24 mg, 0.42 mmol), and iron (III) chloride (4.2 mg, 0.03 mmol) in ethanol (2 mL) and water (2 mL) was heated to 100ºC while stirring. After 2.5 hours, additional iron powder (24 mg, 0.42 mmol) was added, and the mixture continued stirring for 1.5 hours. The mixture was returned to room temperature and filtered through a pad of celite. The crude product was purified by ISCO Flash column chromatography (DCM:MeOH, 0-40%) to afford 62 mg of the desired product (47%). ^1^H NMR (500 MHz, DMSO-*d*_6_): δ 10.10 (s, 1H), 7.02 (d, *J* = 8.6 Hz, 1H), 6.98 (d, *J* = 8.5 Hz, 1H), 6.91 (d, *J* = 2.7 Hz, 1H), 6.84 (d, *J* = 2.7 Hz, 1H), 6.73-6.70 (m, 2H), 5.18 (s, 2H), 3.72 (s, 3H); ^13^C NMR (125 MHz, DMSO-*d*_6_): δ 166.1, 156.7, 152.3, 149.3, 146.2, 125.9, 124.4, 122.1, 120.8, 118.8, 114.6, 111.0, 106.3, 55.5; HRMS: (ESI+) m/z calculated for C_14_H_12_N_2_O_3_ [M+H]^+^: 257.0921, found: 257.0925.

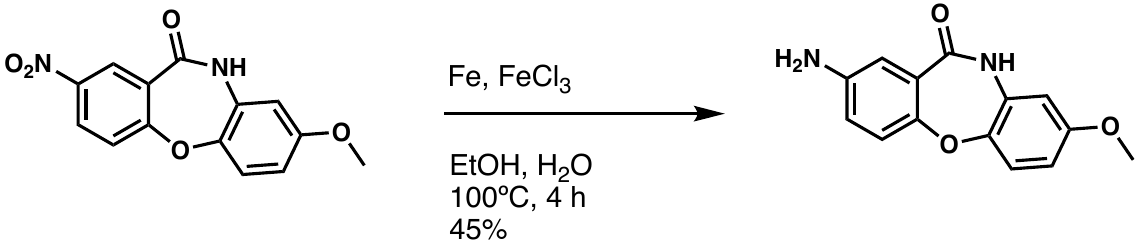

In a round bottom flask, a solution of 8-methoxy-2-nitrodibenz[b,f][1,4]oxazepin-11(10H)-one (150 mg, 0.52 mmol), iron (24 mg, 0.42 mmol), and iron (III) chloride (4.2 mg, 0.03 mmol) in ethanol (2 mL) and water (2 mL) was heated to 100ºC while stirring. After 2.5 hours, additional iron powder (24 mg, 0.42 mmol) was added, and the mixture continued stirring for 1.5 hours. The mixture was returned to room temperature and filtered through a pad of celite. The crude product was purified by ISCO Flash column chromatography (DCM:MeOH, 0-40%) to afford 60 mg of the desired product (45%). ^1^H NMR (500 MHz, DMSO-*d*_6_): δ 10.21 (s, 1H), 7.13 (d, *J* = 8.8 Hz, 1H), 6.95 (d, *J* = 8.5 Hz, 1H), 6.91 (d, *J* = 2.9 Hz, 1H), 6.72 (dd, *J* = 8.8 Hz, 1H), 6.67 (d, *J* = 2.9 Hz, 1H), 6.64 (dd, *J* = 8.8 Hz, 1H), 5.16 (s, 2H), 3.69 (s, 3H); ^13^C NMR (125 MHz, DMSO-*d*_6_): δ 166.5, 156.3, 149.6, 146.0, 145.0, 132.0, 125.7, 121.3, 120.6, 119.0, 114.6, 109.6, 106.6, 55.5; HRMS: (ESI+) m/z calculated for C_14_H_12_N_2_O_3_ [M+H]^+^: 257.0921, found: 257.0919.

**8.**

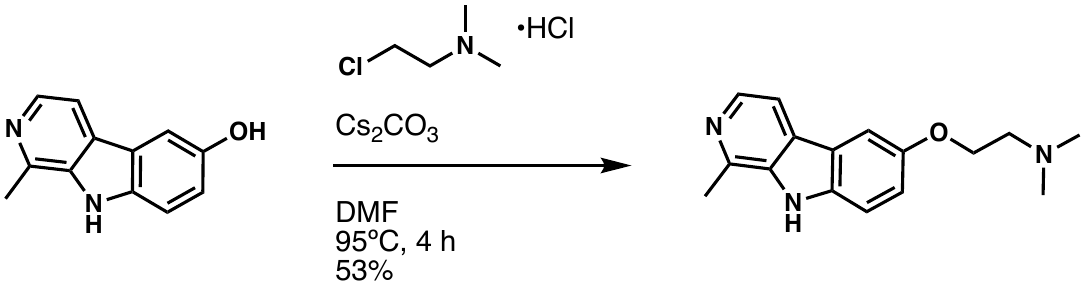

To a solution of harmol (25 mg, 0.126 mmol) in DMF (2.5 mL) was added 2-chloro-N,N-dimethyl-ethanamine hydrochloride (27 mg, 0.189 mmol) and cesium carbonate (62 mg, 0.189 mmol). The mixture was heated to reflux and stirred for four hours. After cooling to room temperature, the reaction was diluted with water (10 mL), and the mixture was extracted with dichloromethane (3 x 10 mL). The combined organic layer was then washed with brine (10 mL) and dried with Na_2_SO_4_. The crude product was concentrated down onto silica gel and purified by ISCO Flash Column Chromatography (DCM:MeOH, 5-40%) to afford 18 mg of yellowish brown solid (53%). ^1^H NMR (500 MHz, DMSO-*d*_6_): δ 11.80 (s, 1H), 8.15 (d, *J* = 5.1 Hz, 1H), 8.07 (d, *J* = 9.0 Hz, 1H), 7.83 (d, *J* = 4.8 Hz, 1H), 7.08 (s, 1H), 6.87 (d, *J* = 9.0 Hz, 1H), 4.34 (t, *J* = 5.0 Hz, 2H), 3.16 (t, *J* = 5.6 Hz, 2H), 2.74 (s, 3H), 2.57 (s, 6H); ^13^C NMR (125 MHz, DMSO-*d*_6_): δ 158.7, 142.0, 141.3, 137.3, 134.6, 127.3, 122.7, 115.1, 112.0, 109.4, 95.7, 64.3, 56.3, 44.0, 20.3; HRMS: (ESI+) m/z calculated for C_16_H_19_N_3_O [M+H]^+^: 270.1601, found: 270.1601.

**NMR Spectra**

**1**

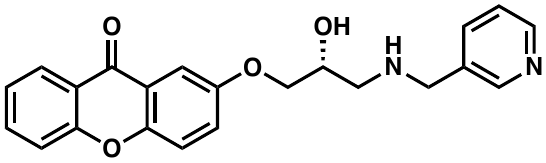

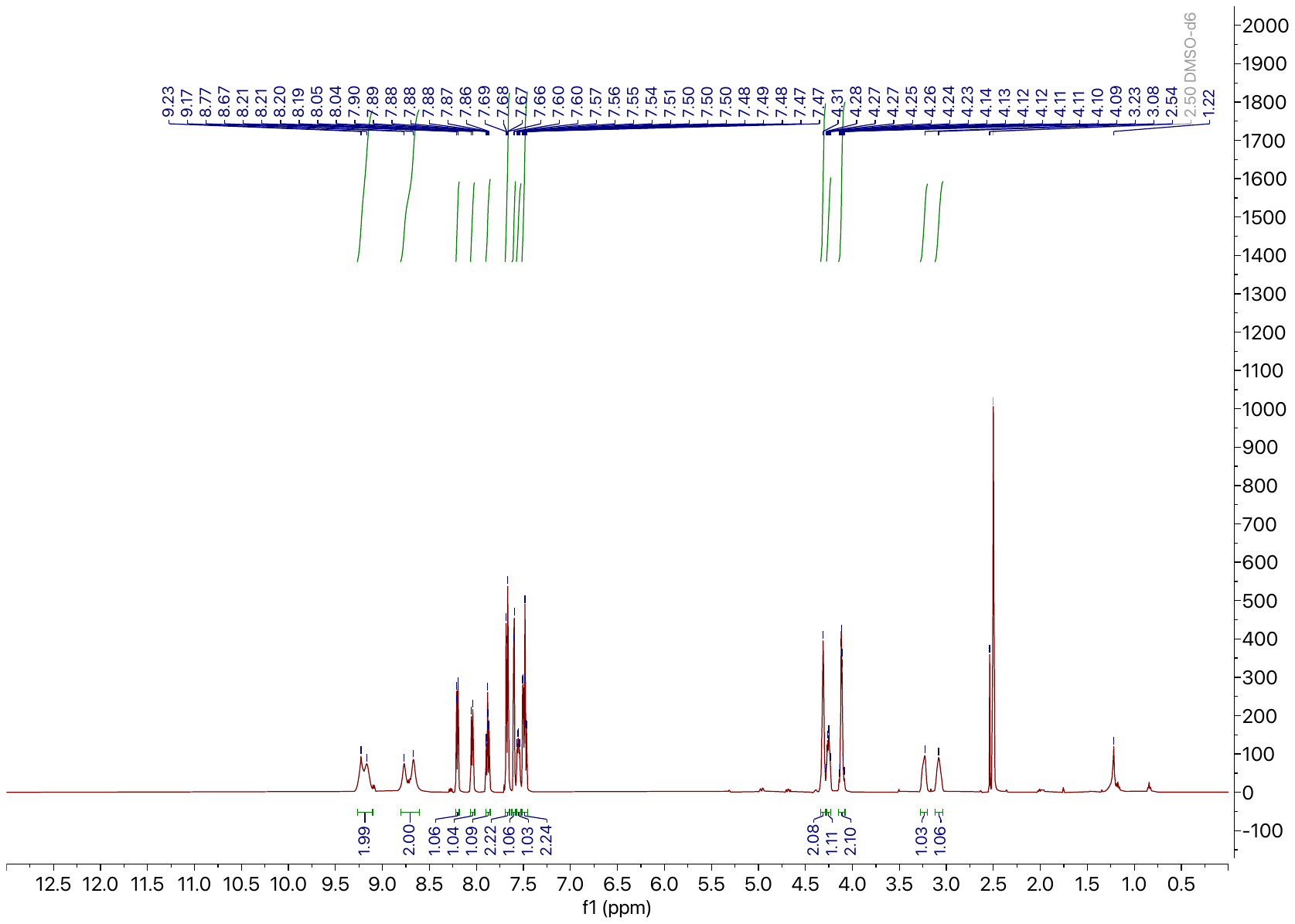

**^1^H NMR.** 500 MHz, DMSO-*d*_6_

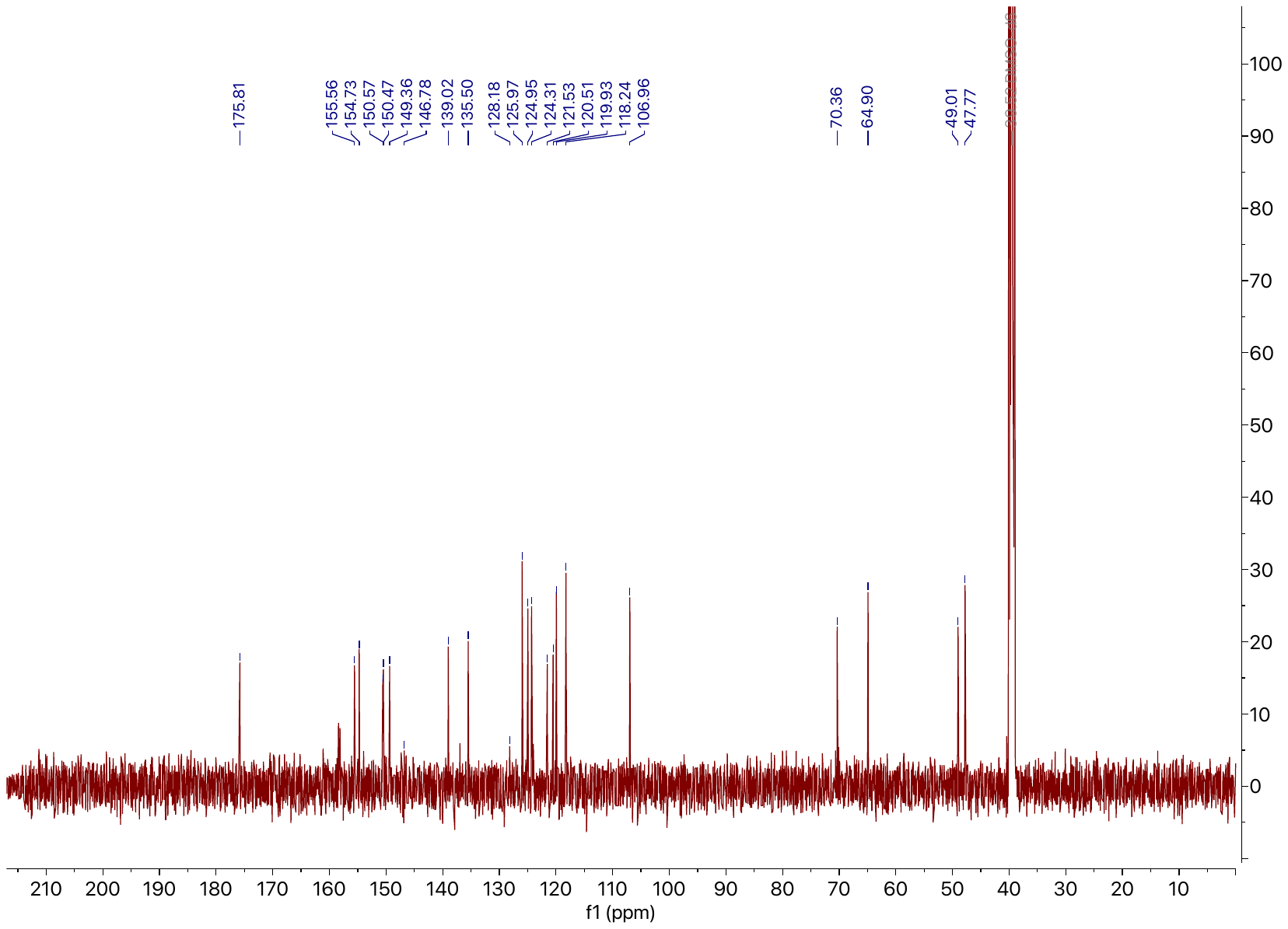

**^13^C NMR.** 125 MHz, DMSO-*d*_6_

**2**

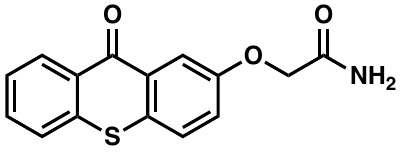

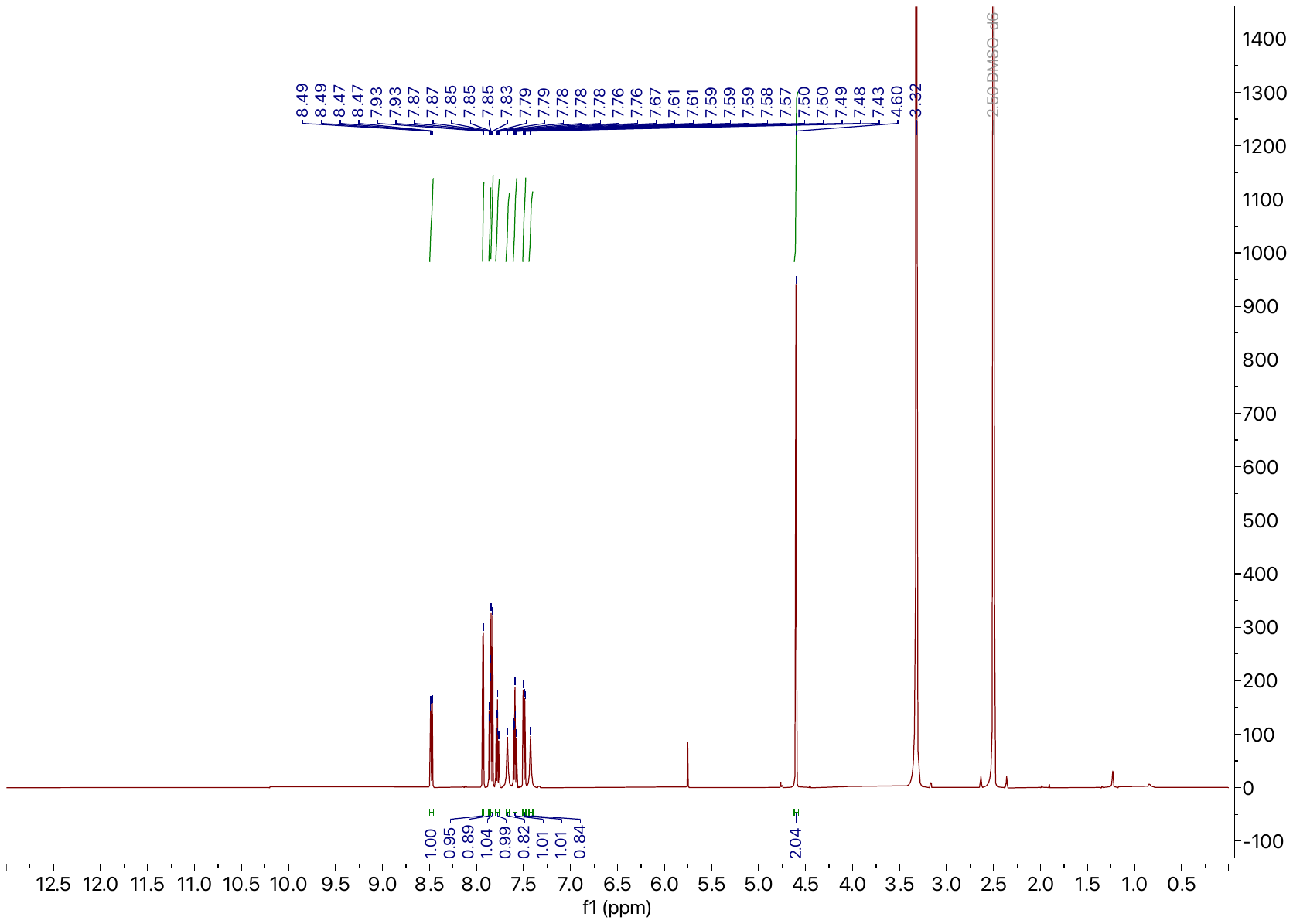

**^1^H NMR.** 500 MHz, DMSO-*d*_6_

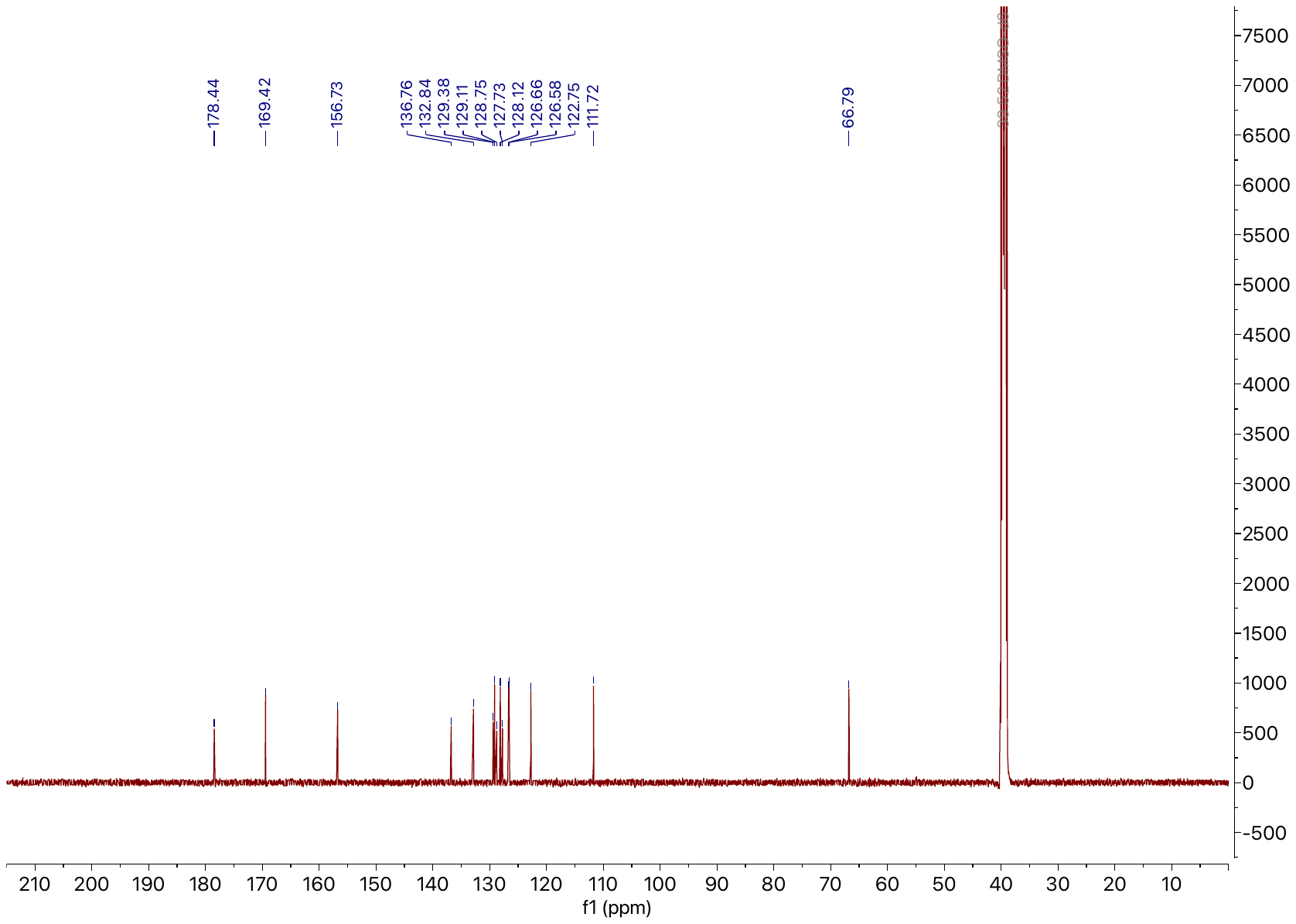

**^13^C NMR.** 125 MHz, DMSO-*d*_6_

**3-1**

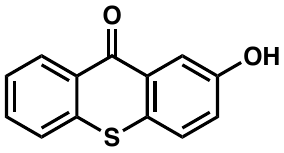

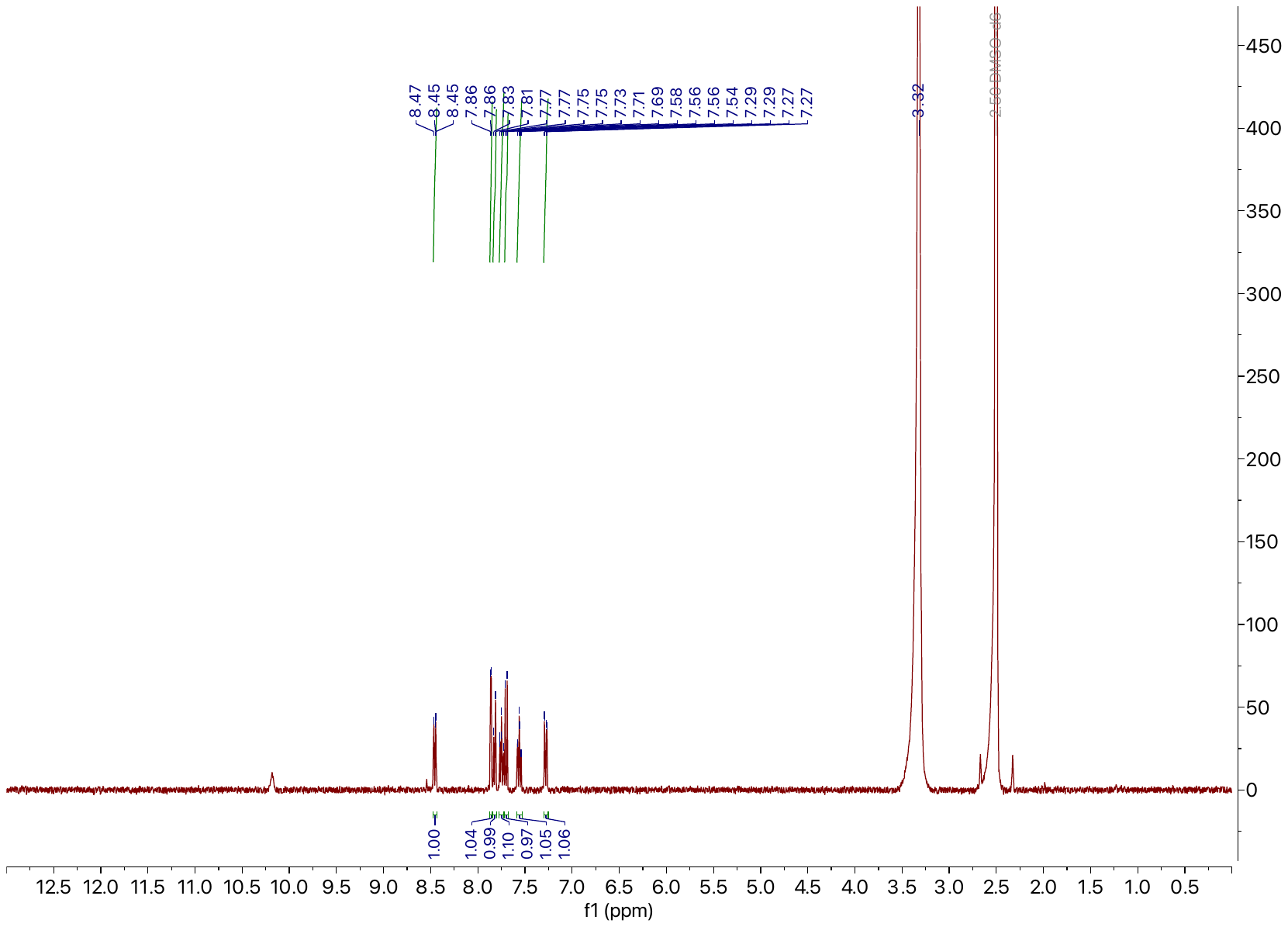

**^1^H NMR.** 500 MHz, DMSO-*d*_6_

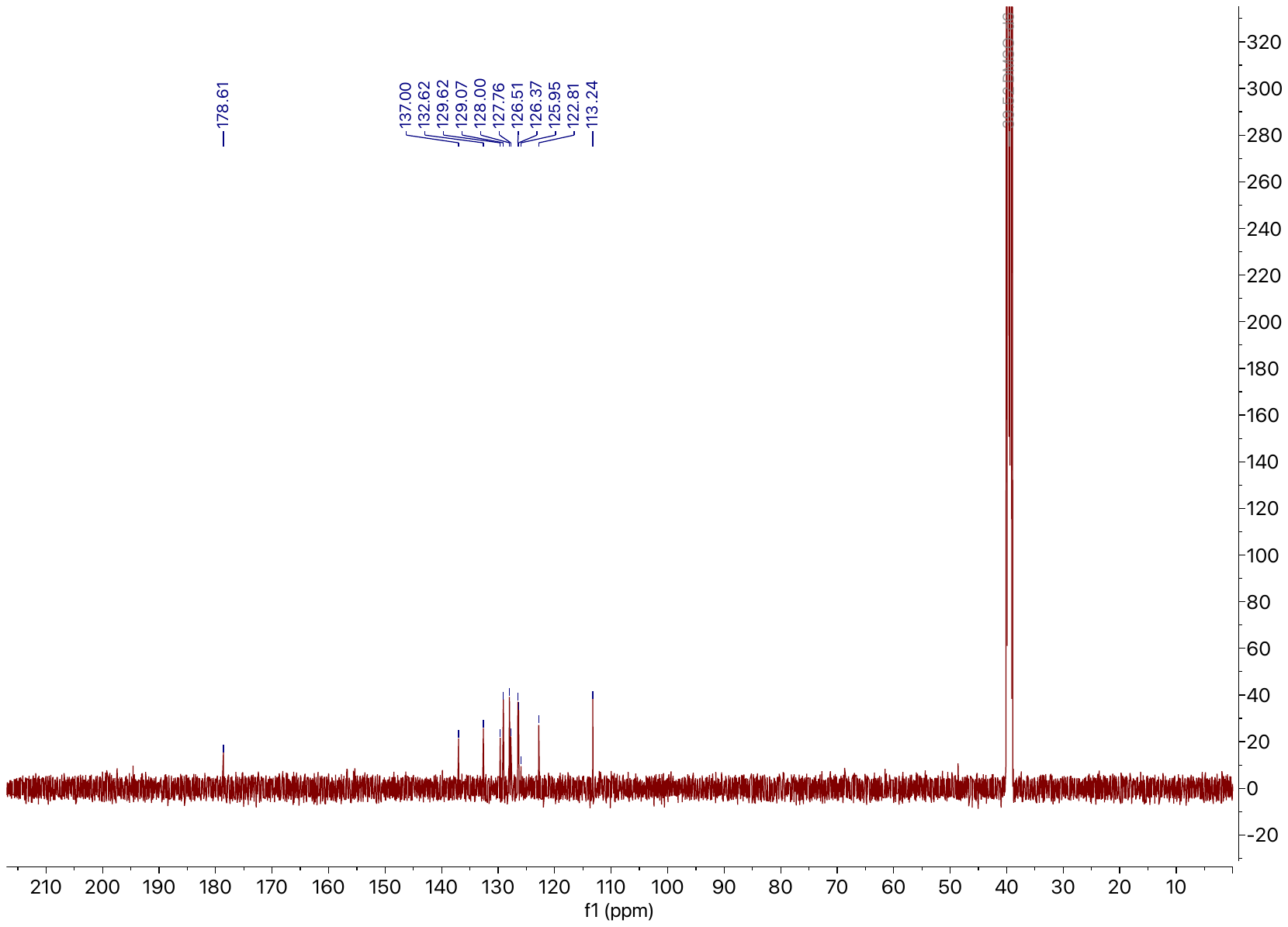

**^13^C NMR.** 125 MHz, DMSO-*d*_6_

**3-2**

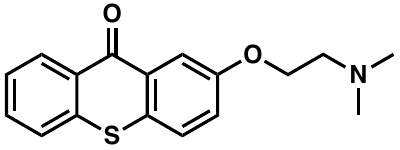

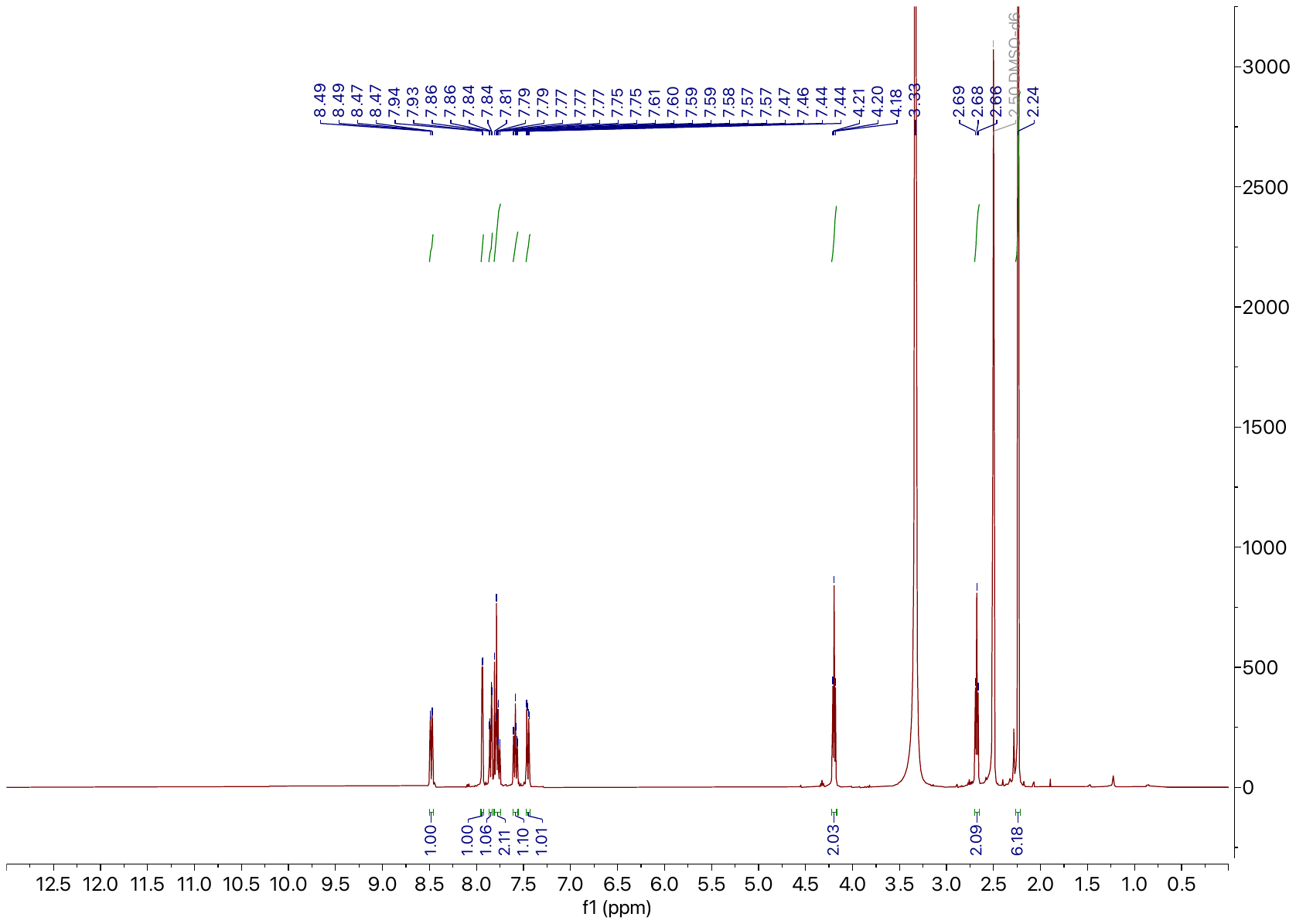

**^1^H NMR.** 500 MHz, DMSO-*d*_6_

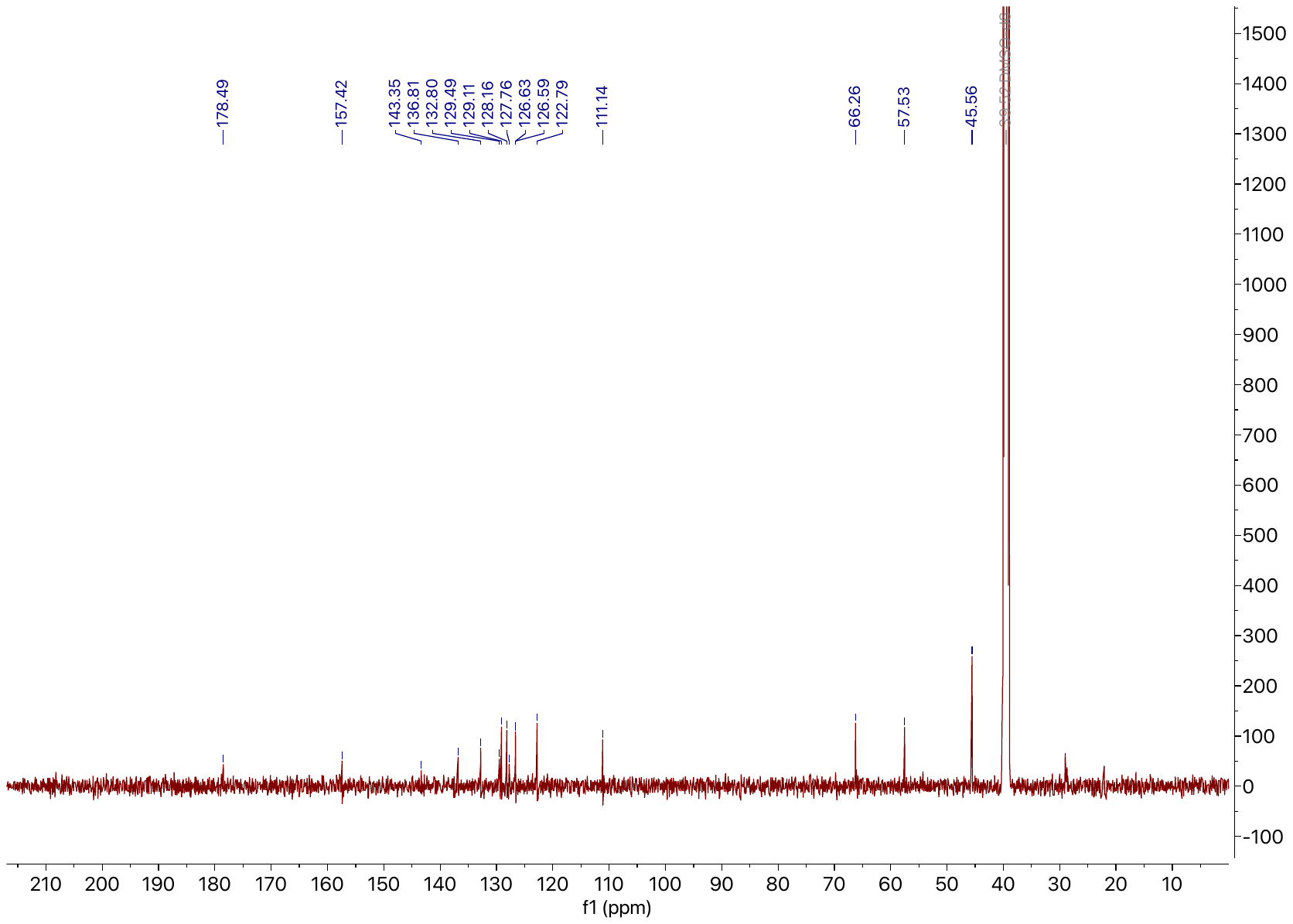

**^13^C NMR.** 125 MHz, DMSO-*d*_6_

**4**

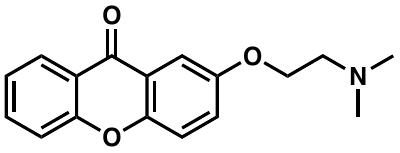

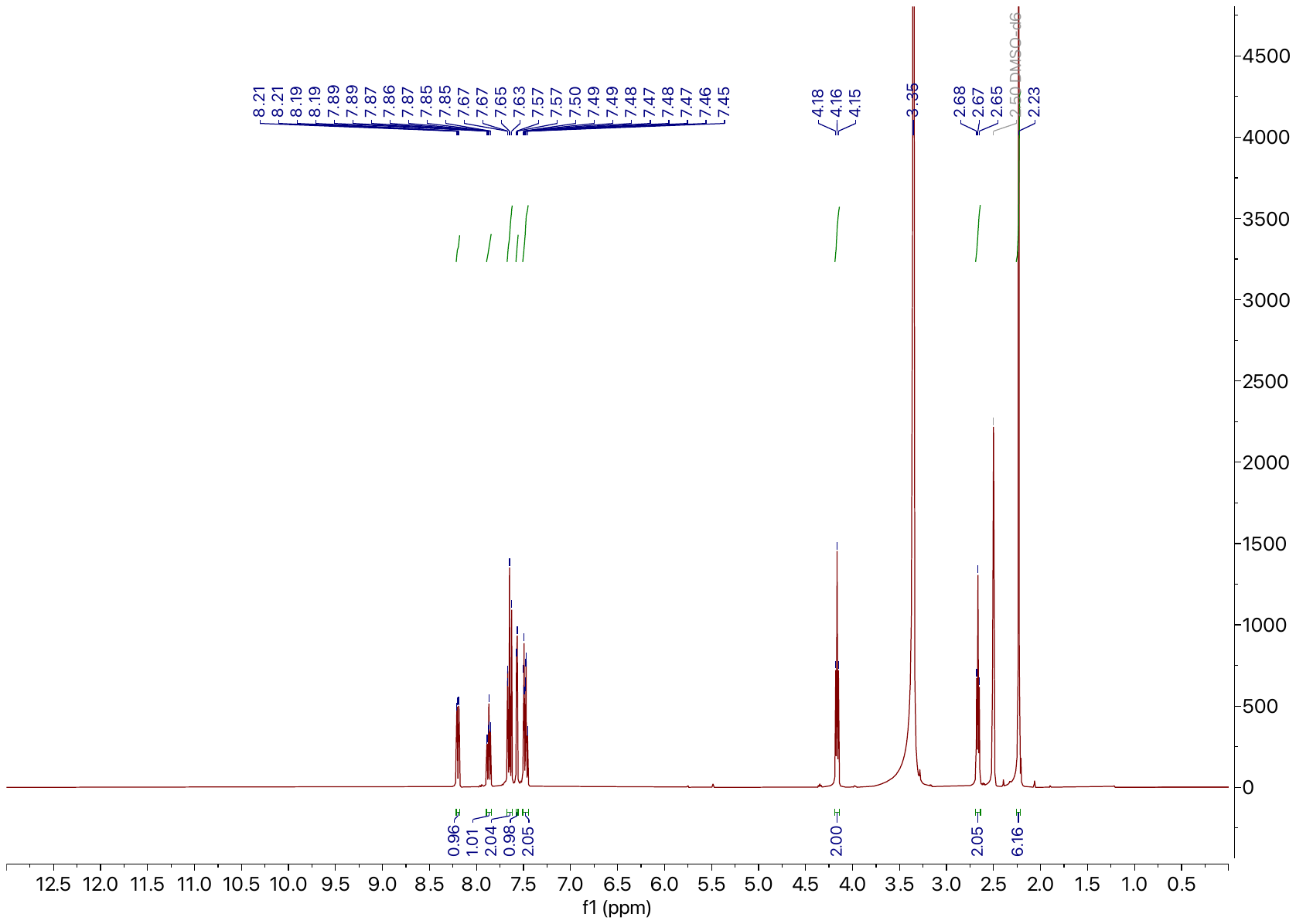

**^1^H NMR.** 500 MHz, DMSO-*d*_6_

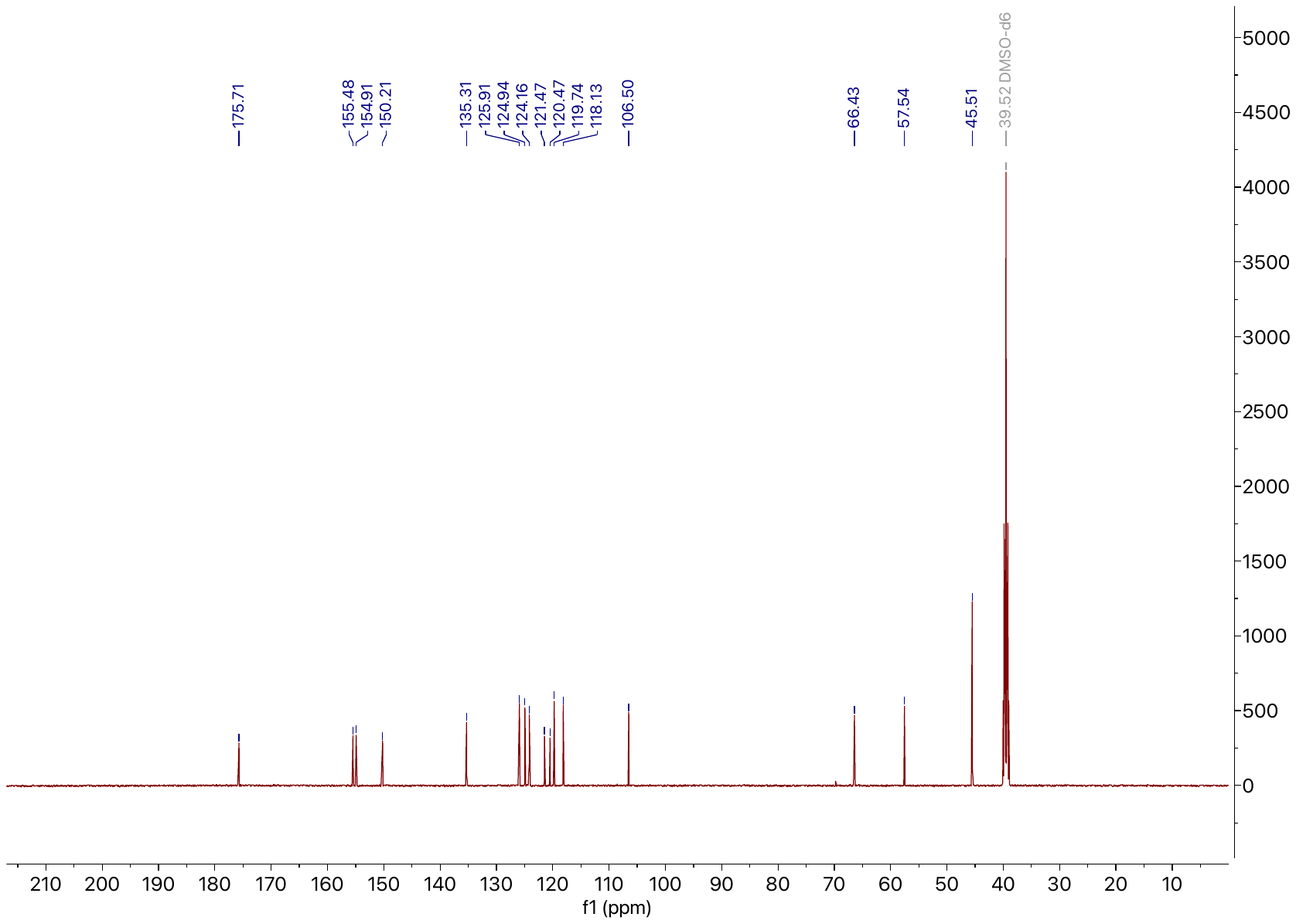

**^13^C NMR.** 125 MHz, DMSO-*d*_6_

**5**

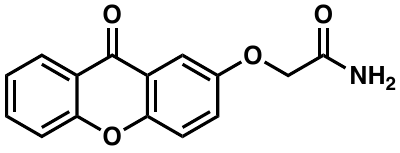

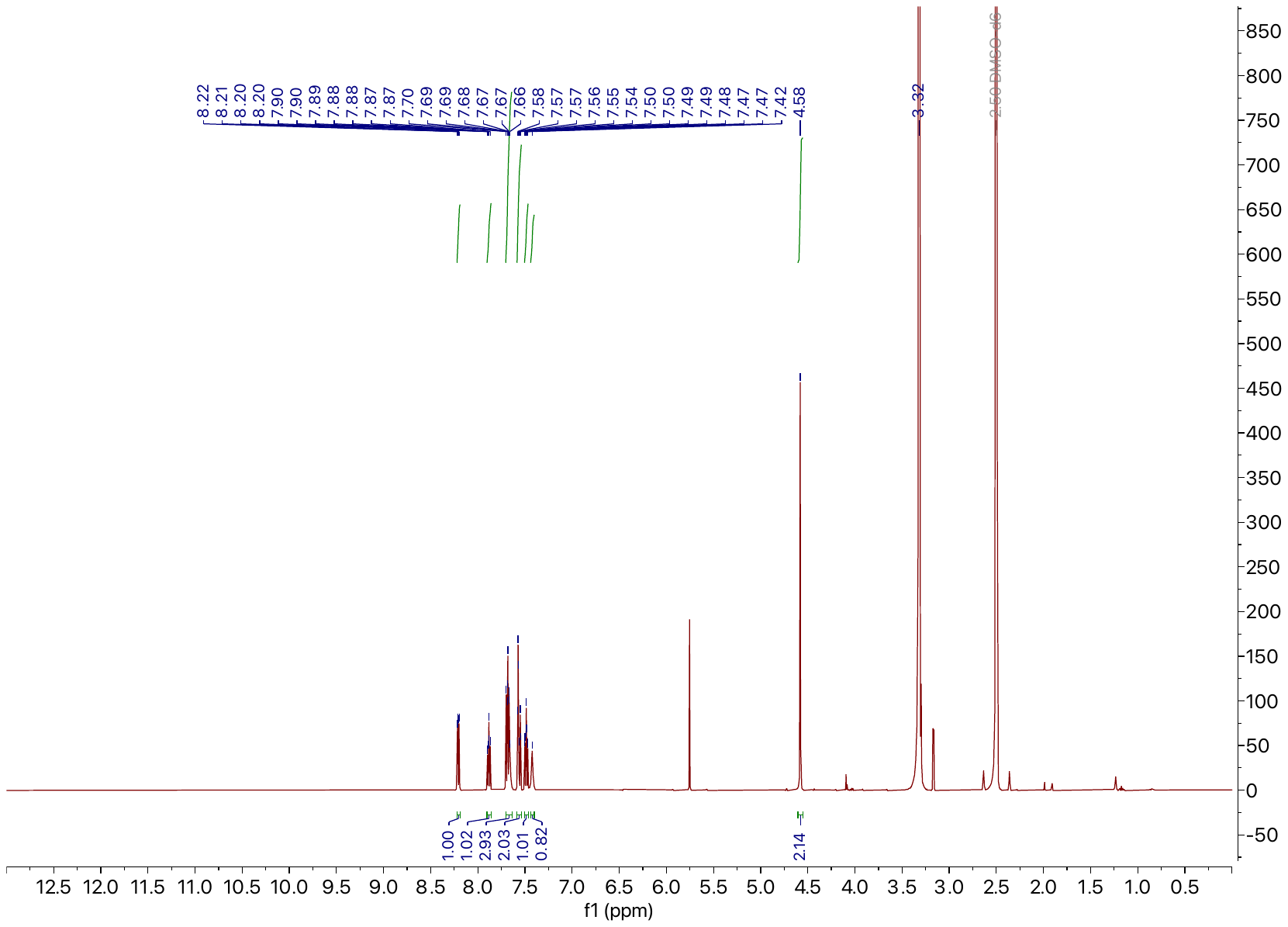

**^1^H NMR.** 500 MHz, DMSO-*d*_6_

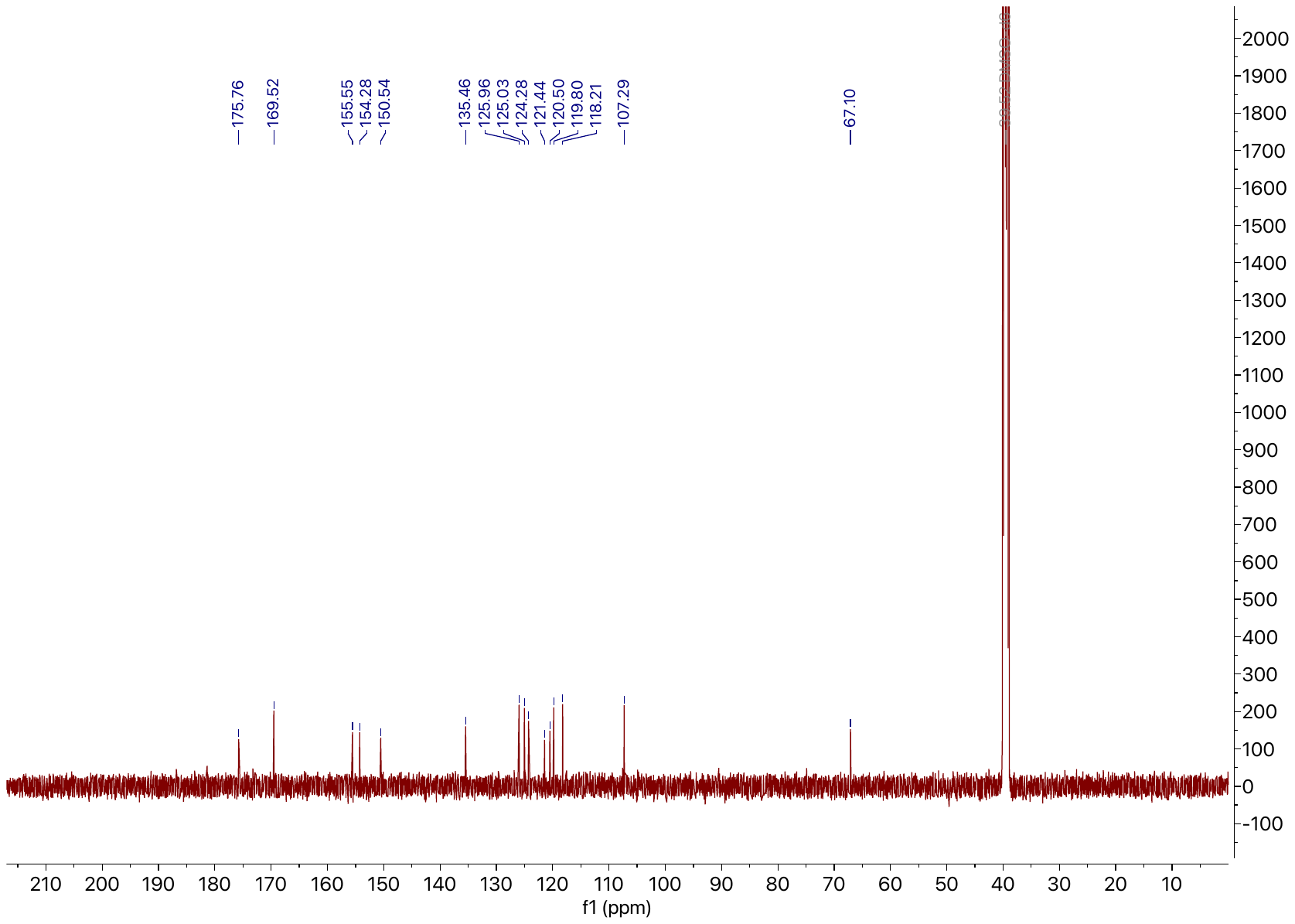

**^13^C NMR.** 125 MHz, DMSO-*d*_6_

**6**

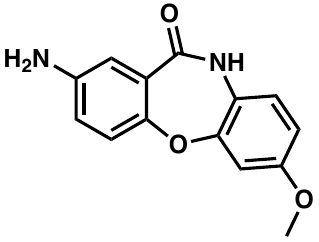

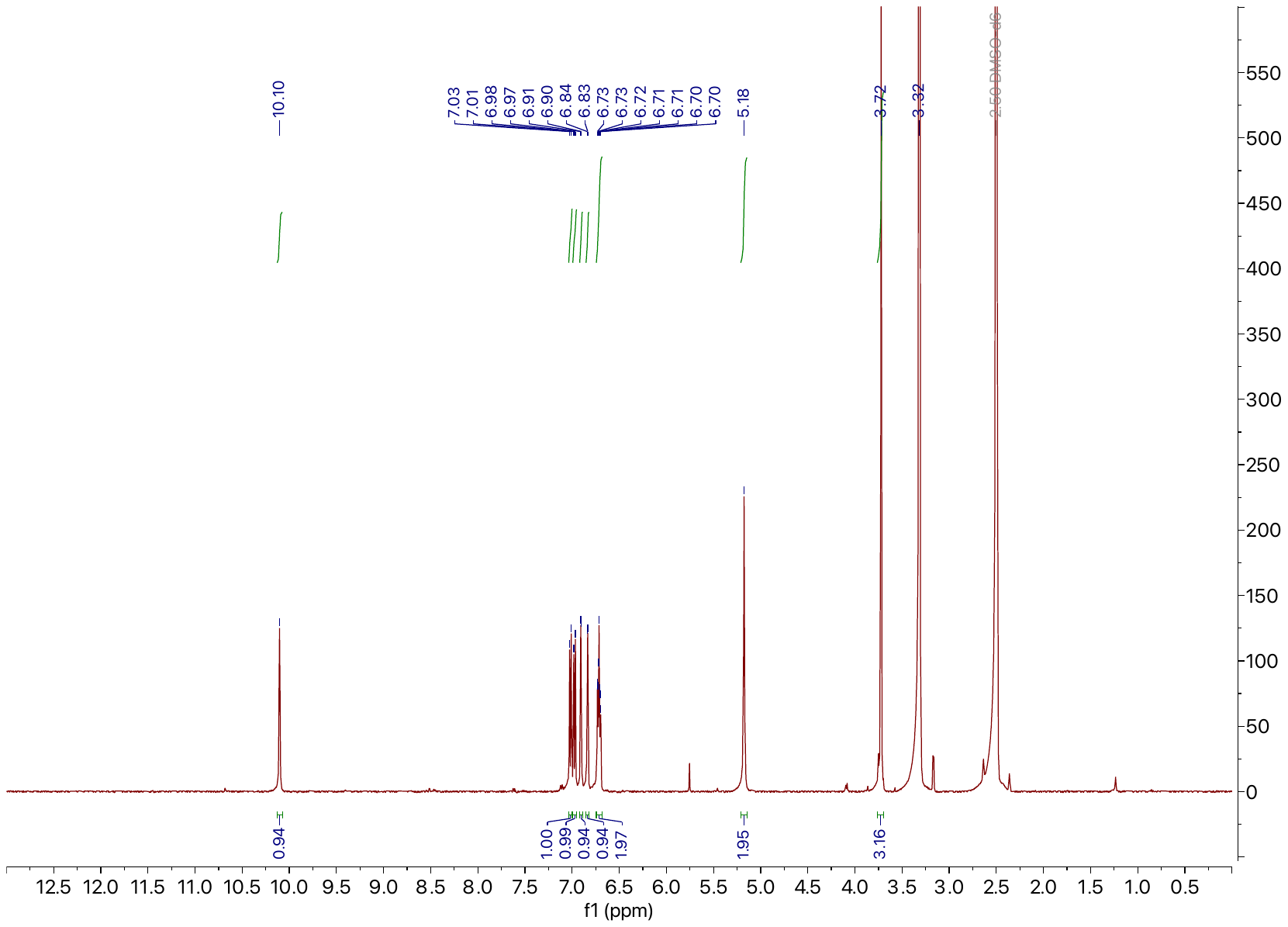

**^1^H NMR.** 500 MHz, DMSO-*d*_6_

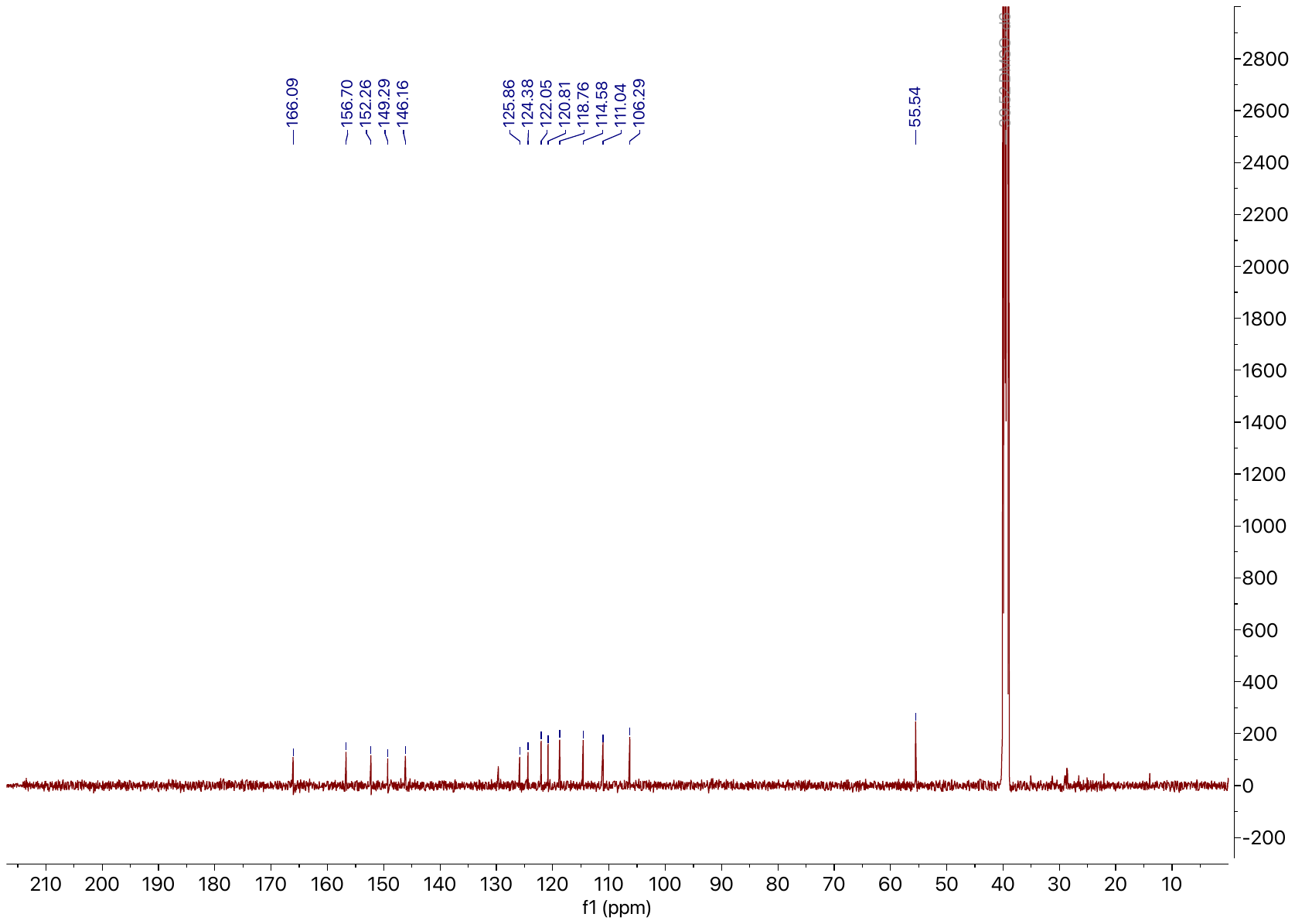

**^13^C NMR.** 125 MHz, DMSO-*d*_6_

**7**

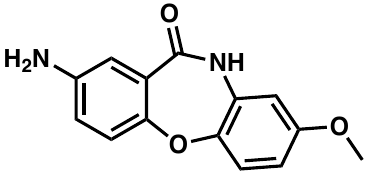

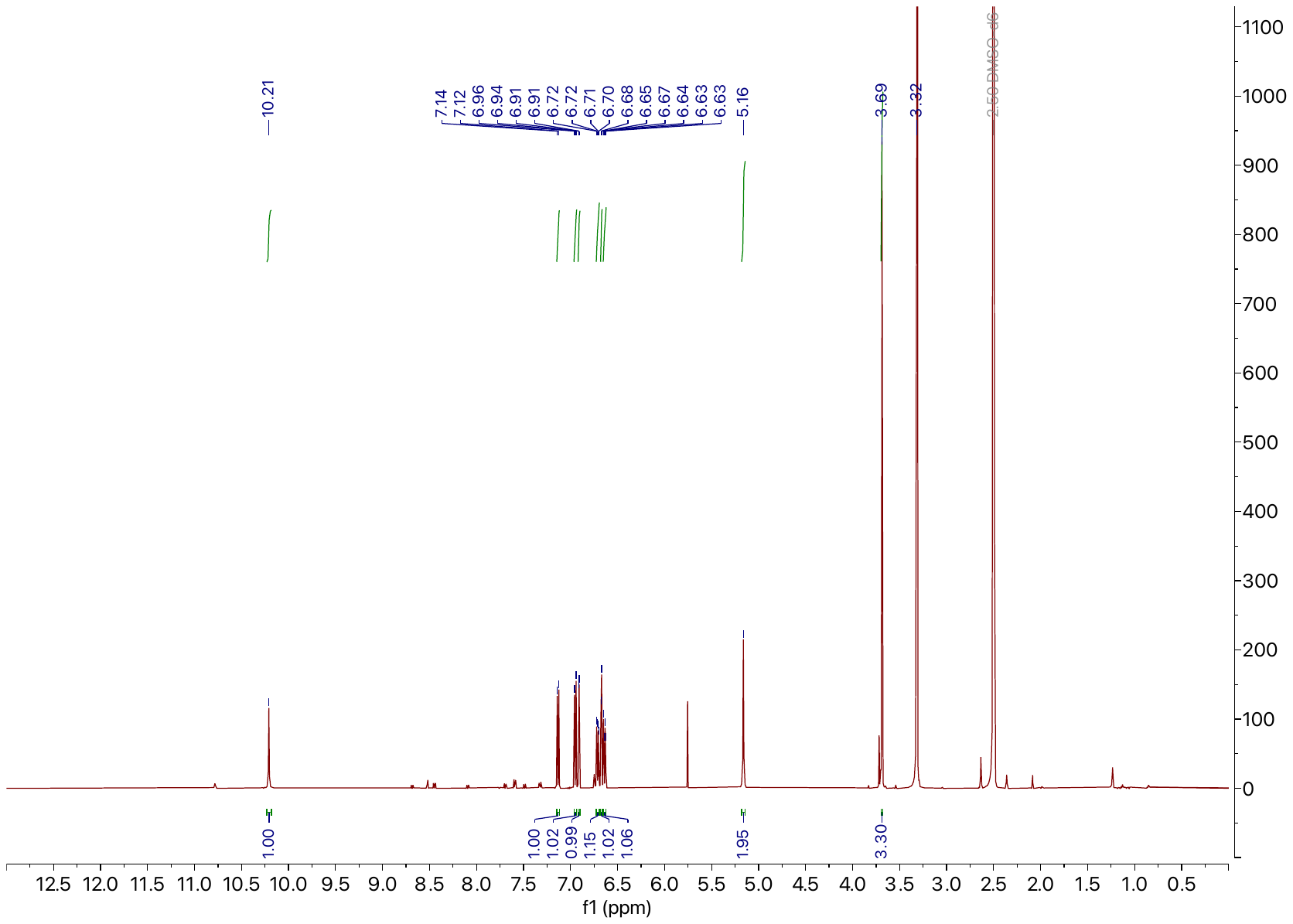

**^1^H NMR.** 500 MHz, DMSO-*d*_6_

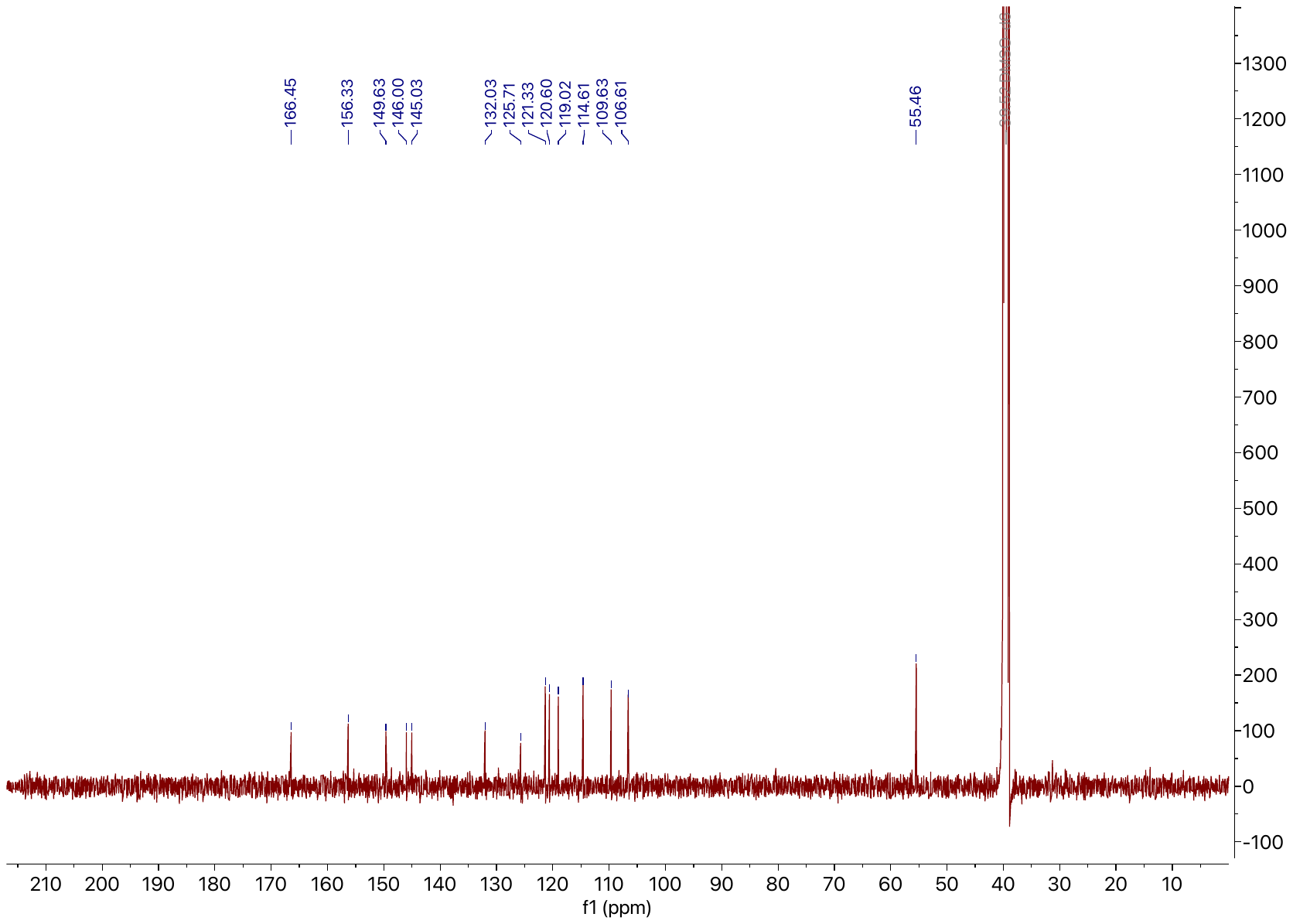

**^13^C NMR.** 125 MHz, DMSO-*d*_6_
**8**

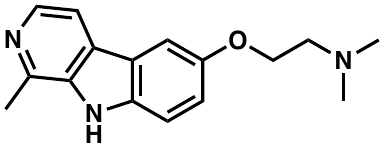

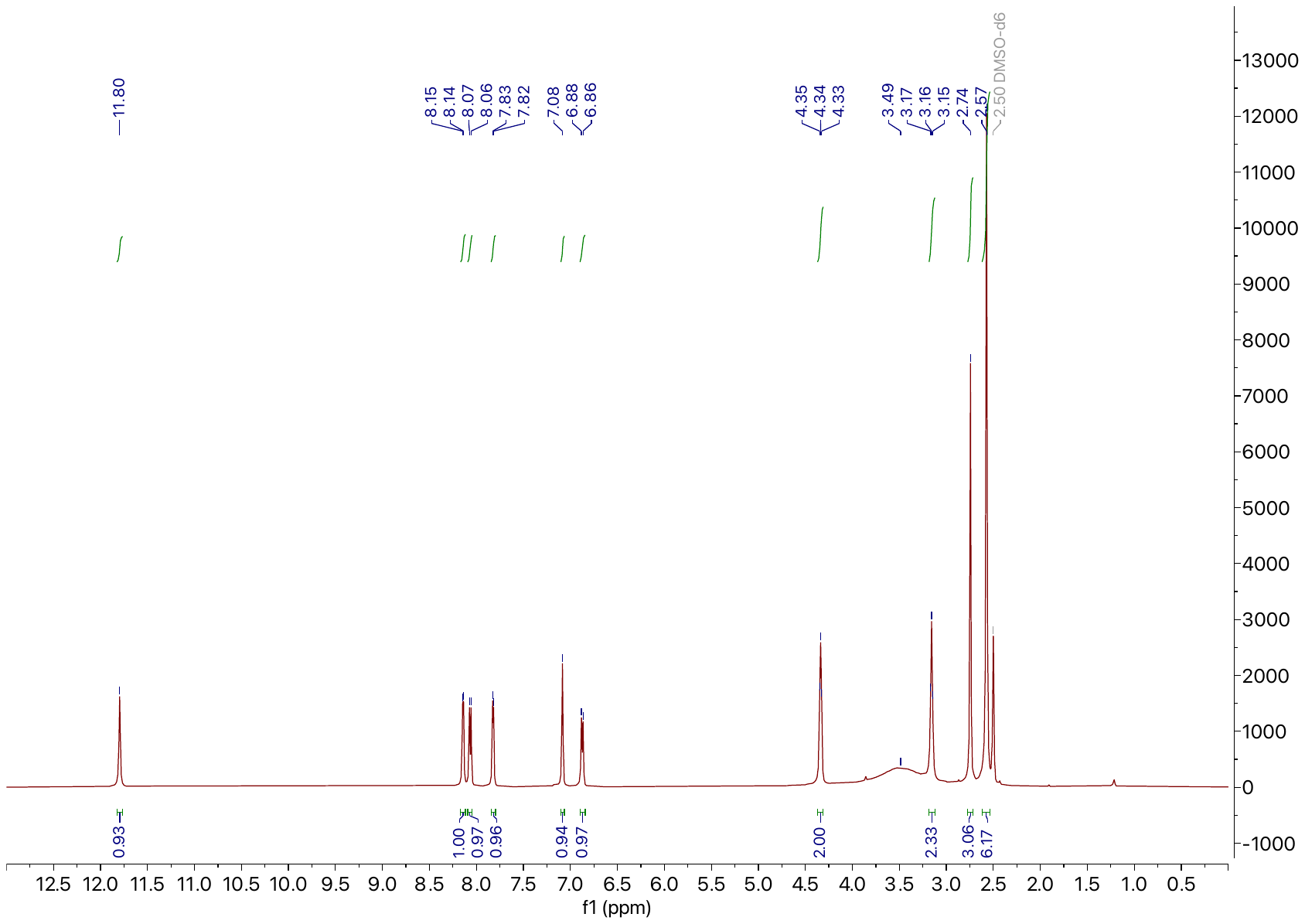

**^1^H NMR.** 500 MHz, DMSO-*d*_6_

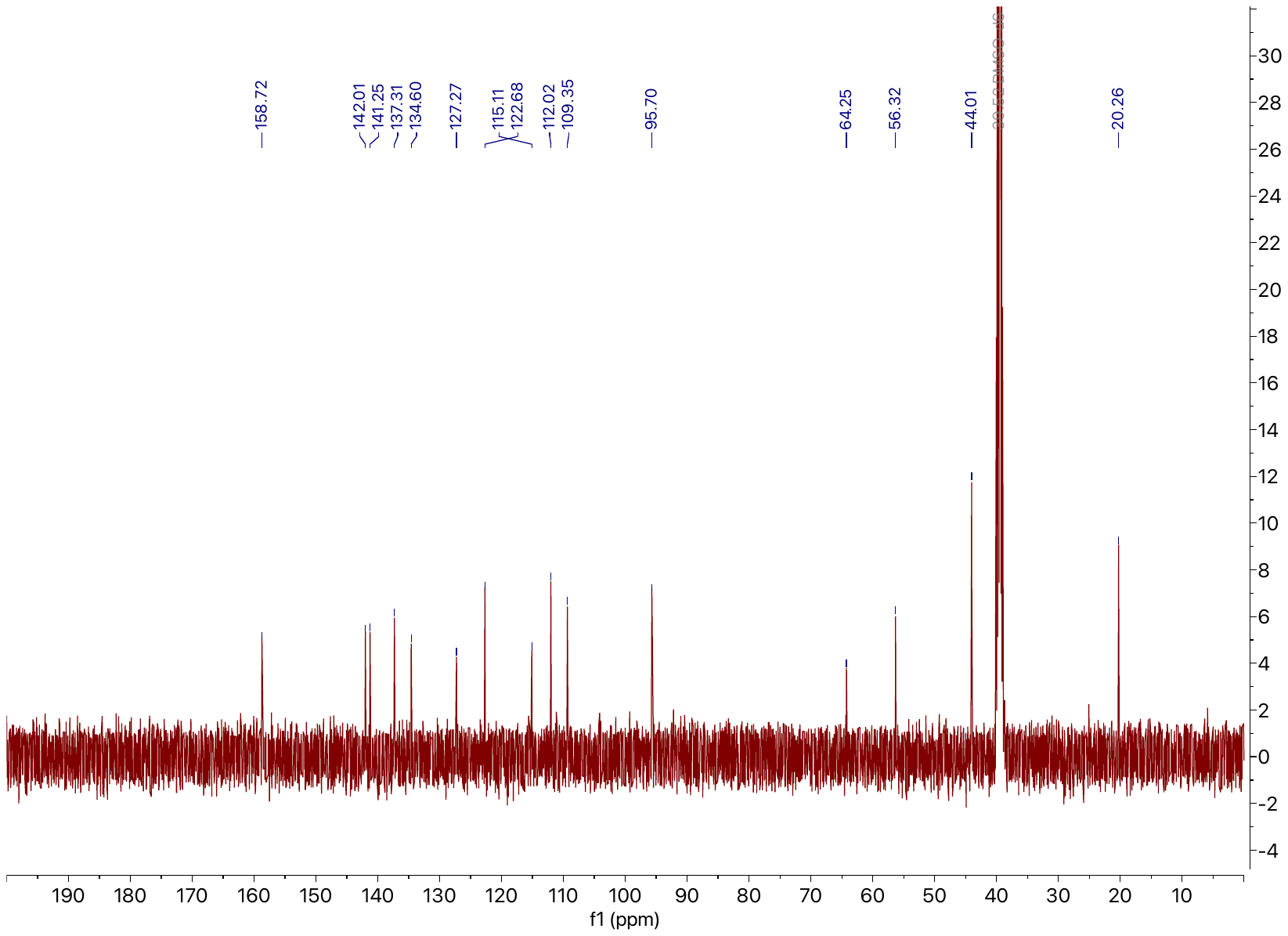

**^13^C NMR.** 125 MHz, DMSO-*d*_6_
